# A CRISPR-Cas9–engineered mouse model for GPI-anchor deficiency mirrors human phenotypes and exhibits hippocampal synaptic dysfunctions

**DOI:** 10.1101/2020.04.20.050591

**Authors:** Miguel Rodríguez de los Santos, Marion Rivalan, Friederike S. David, Alexander Stumpf, Julika Pitsch, Despina Tsortouktzidis, Laura Moreno Velasquez, Anne Voigt, Karl Schilling, Daniele Mattei, Melissa Long, Guido Vogt, Alexej Knaus, Björn Fischer-Zirnsak, Lars Wittler, Bernd Timmermann, Peter N. Robinson, Denise Horn, Stefan Mundlos, Uwe Kornak, Albert J. Becker, Dietmar Schmitz, York Winter, Peter M. Krawitz

**Author notes:** Corresponding author: Professor Dr. med. Dipl. phys. Peter Krawitz, Institute for Genomic Statistics and Bioinformatics, University Hospital Bonn, Venusberg, Campus 1, 53127 Bonn, Germany.

## Abstract

Pathogenic germline mutations in *PIGV* lead to glycosylphosphatidylinositol biosynthesis deficiency (GPIBD). Individuals with pathogenic biallelic mutations in genes of the glycosylphosphatidylinositol (GPI) anchor pathway exhibit cognitive impairments, motor delay, and often epilepsy. Thus far, the pathophysiology underlying the disease remains unclear, and suitable rodent models that mirror all symptoms observed in human patients have not been available. Therefore, we used CRISPR-Cas9 to introduce the most prevalent hypomorphic missense mutation in European patients, *Pigv:c*.1022C>A (p.A341E), at a site that is conserved in mice. Mirroring the human pathology, mutant *Pigv*^341E^ mice exhibited deficits in motor coordination, cognitive impairments, and alterations in sociability and sleep patterns, as well as increased seizure susceptibility. Furthermore, immunohistochemistry revealed reduced synaptophysin immunoreactivity in *Pigv*^341E^ mice, and electrophysiology recordings showed decreased hippocampal synaptic transmission that could underlie impaired memory formation. In single-cell RNA sequencing, *Pigv*^341E^-hippocampal cells exhibited changes in gene expression, most prominently in a subtype of microglia and subicular neurons. A significant reduction in *Abl1* transcript levels in several cell clusters suggested a link to the signaling pathway of GPI-anchored ephrins. We also observed elevated levels of *Hdc* transcripts, which might affect histamine metabolism with consequences for circadian rhythm. This new mouse model will not only open the doors to further investigation into the pathophysiology of GPIBD, but will also deepen our understanding of the role of GPI-anchor–related pathways in brain development.

**Significance statement:** Inherited GPI-anchor biosynthesis deficiencies (IGDs) explain many cases of syndromic intellectual disability. Although diagnostic methods are improving, the pathophysiology underlying the disease remains unclear. Furthermore, we lack rodent models suitable for characterizing cognitive and social disabilities. To address this issue, we generated the first viable mouse model for an IGD that mirrors the condition in human patients with a behavioral phenotype and susceptibility to epilepsy. Using the new model, we obtained neurological insights such as deficits in synaptic transmission that will facilitate understanding of the pathophysiology of IGDs.

## Introduction

The glycosylphosphatidylinositol (GPI) anchor is essential for connecting a remarkable number of proteins (GPI-linked proteins) to the cell membrane. GPI-linked proteins are essential for signal transduction, cell–cell adhesion, axonal outgrowth, synapse formation, and plasticity, as well as for regulation of the complement system (1,2). Paroxysmal nocturnal hemoglobinuria (PNH) was the first disorder to be characterized as a GPI-anchor biosynthesis deficiency (GPIBD) (3). However, PNH is exceptional in two regards: First, it is the only GPIBD that is acquired and it is due to somatic mutations that cause complete loss of function. In inherited GPIBDs, also referred to as inherited GPI-anchor biosynthesis deficiencies (IGDs), residual GPI-anchor synthesis and maturation activities persist. Second, the prevalence of inherited GPIBDs is at least 10-fold higher than that of PNH. To date, recessive phenotypes have been reported for 21 genes of the GPI-anchor pathway. Bellai-Dussault discussed the clinical variability in detail for the first 19 GPIBDs (4). However, most patients, including recently described cases due to GPIBD20 and GPIBD21, exhibit intellectual disability, psychomotor delay, and epilepsy (5,6). Furthermore, due to the residual GPI-anchor synthesis and maturation, patient-derived fibroblasts have a reduced number of GPI-linked proteins on the cell surface (7).

Prior to the discovery of IGD, mouse models of GPI anchor deficiency, which mainly employed chimeric and conditional knockouts in which GPI-anchor biosynthesis was abolished in specific tissues, demonstrated that a complete loss of GPI anchors is embryonic lethal (8–12). Interestingly, the resultant phenotypes were often still so severe that the mutant mice died early, suggesting essential functions of GPI-anchor proteins in the skin, development of white matter, and dendritic arborization of Purkinje cells in the cerebellum (8,10). In recent years, mice with constitutional GPIBDs were identified in mutation screens; these animals were viable probably because the mutations were only hypomorphic or affected isoforms that are limited to certain tissues, and therefore only explain some aspects of most inherited GPIBDs (13,14). Lukacs et al. showed that a missense mutation in *Pgap2*, p.(M1V), compromised transcription of this gene particularly in neural crest cells, resulting in a craniofacial phenotype in mutant mice. In contrast to most other genes involved in GPI-anchor biosynthesis and maturation, *Pgap2* has a tissue-specific expression pattern that changes over embryonic development, and the existence of multiple isoforms in humans further complicates phenotypic comparisons between humans and mice. It is likely that the facial abnormalities described in human patients with a similar mutation, p.(M1R), are due to a similar mechanism; however, the other phenotypic features seem to be milder in humans, either because the substituted amino acid is different or because the isoforms resulting in p.(M52R) and p.(M58R) do not exist in mice. In addition, McKean et al. observed a holoprosencephaly-like phenotype in two mouse models with a frame-shift mutation in *Pign* and an in-frame deletion in *Pgap1* (14).

Because most of the existing mouse models die at an early stage due to the severe phenotype, it has not been possible to use these models to characterize cognitive deficits, which represent the main challenge in individuals with IGDs. For this purpose, we used CRISPR-Cas9 to engineer a mouse model with the missense mutation c.1022C>A, p.A341E in *Pigv*, one of the most frequently encountered pathogenic alleles in humans (Krawitz et al. 2010). *PIGV* encodes mannosyl transferase II, which is essential for the attachment of the second mannose to the GPI anchor (16). Due to the residual function of *Pigv*^341E^, mutant mice are viable with a normal life span, making it possible to complete behavioral experiments that test motor, social, and cognitive abilities, and study the brain tissue, as well as cells of these mice.

## Results

Patients with IGDs exhibit a heterogeneous spectrum of symptoms, including neurologic findings, movement disorders, and intellectual disability (7,17–19). In general, IGDs caused by pathogenic mutations in genes that catalyze the early steps of GPI-anchor synthesis, such as *PIGA*, tend to have more severe clinical features such as status epilepticus. In patients with mutations in later steps of synthesis, such as *PIGV*, epilepsies also occur in a substantial proportion of cases; however, they often disappear later in life, and intellectual disability becomes the key clinical feature. In contrast to acquired GPIBDs, such as PNH, all patients with IGDs have some residual function of GPI-anchor synthesis, implying that null mutants are not viable and that at least one hypomorphic allele must be present. When a novel mutation is encountered in a suspected IGD, flow cytometry with two or more different markers serves to confirm a GPIBD. To analyze the effect of *Pigv*^341E^ on a cellular level, we used fluor-proaerolysin (FLAER), which can recognize all GPI-anchored proteins, and CD90, a GPI-linked protein that is highly expressed in human fibroblasts. The mean fluorescence intensity (MFI) of FLAER was reduced in hom-*Pigv*^341E^ mouse embryonic fibroblasts (MEFs) (Fig. 1A). Likewise, CRISPR-Cas9–engineered mouse embryonic stem (mES) cell clones (hom-*Pigv*^341E^, hom-*Pgap3*^107L^) exhibited partial reductions in FLAER and GPI-anchored CD90 (SI *Appendix*, Fig. S1E). Unlike clones with homozygous hypomorphic mutations, the null mES clone [*Pigv* (-/-)] exhibited almost no cell-surface expression of FLAER and CD90 (SI *Appendix*, Fig. S1E). Therefore, we concluded that *Pigv*^341E^ is hypomorphic in mice, as it is in humans.

**Figure 1.**
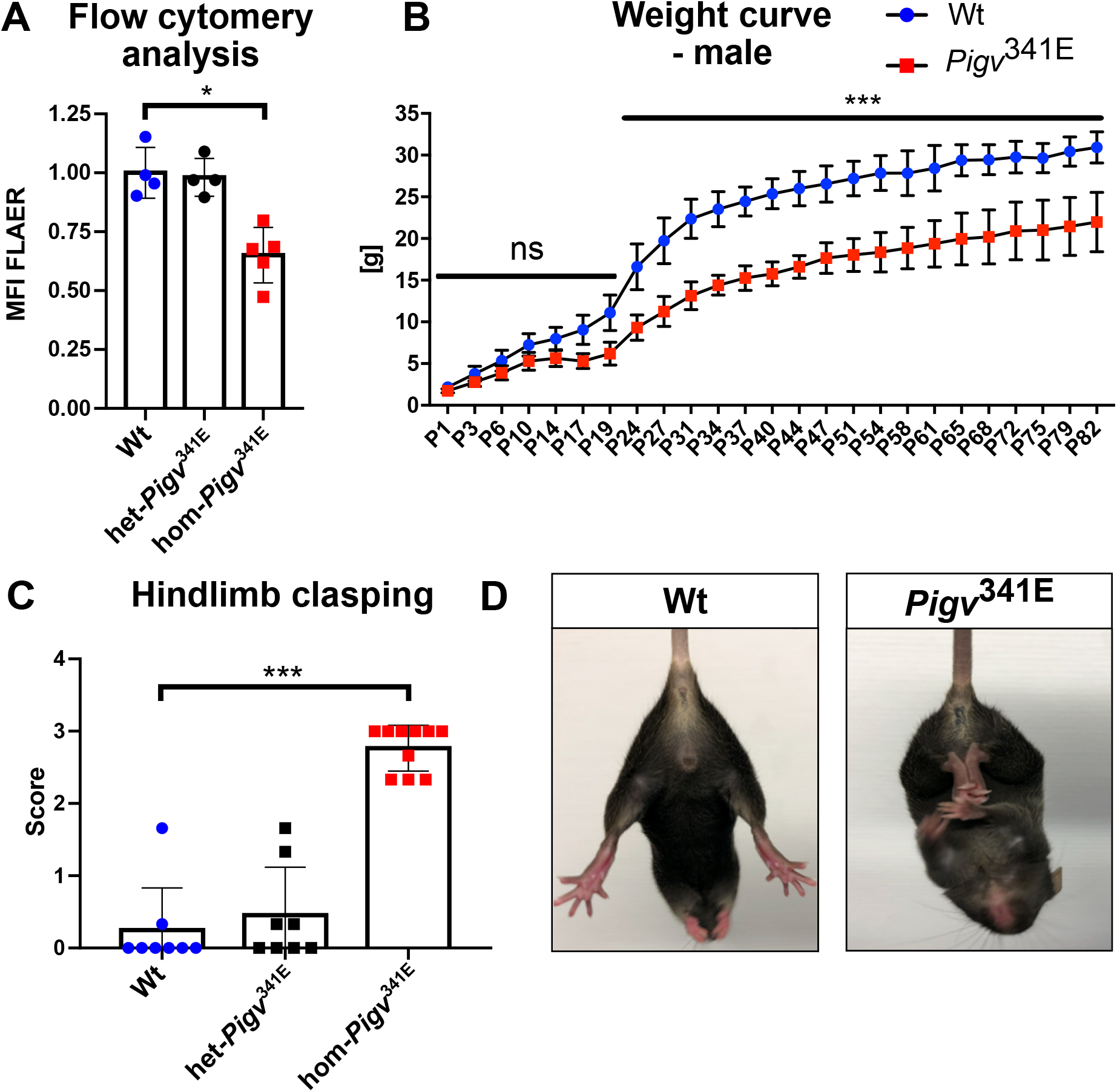
Features of *Pigv*^341E^ mice. (A) Flow cytometry analysis of hom-*Pigv*^341E^ mouse embryonic fibroblasts (MEFs) isolated from embryos (E13,5) revealed a reduced mean fluorescence intensity (MFI) of fluor-proaerolysin (FLAER). MFI was normalized against the wild type. (B) Male *Pigv*^341E^ mice had a reduced weight on postnatal days (P) 1–82. (C) Hom-*Pigv*^341E^ mice exhibited hindlimb clasping behavior. (D) Representative posture of hindlimb clasping behavior in hom-*Pigv*^341E^ mice. By contrast, wild-type mice spread their hindlimbs when picked up by their tail. Hom-*Pigv*^341E^=homozygous for Pigv p.Ala341Glu; het-*Pigv*^341E^=heterozygous for *Pigv* p.Ala341Glu; wt=wild type. Animals used for the weight curve: wt(male=3), hom(male=4). Animals used for the hindlimb clasping test were 6 weeks old: wt(female n=3, male n=5), het-*Pigv*^341E^(female n=4, male n=4), hom-*Pigv*^341E^(female n=4, male= n=6). Data from the weight curve were analyzed by two-way analysis of variance (ANOVA) followed by Bonferroni’s multiple comparisons test. The data from flow cytometry and the hindlimb clasping test were analyzed by non-parametric t-test (Mann–Whitney). *p < 0.05, **p < 0.01, ***p < 0.001.

The results section is structured as follows: We will start with a description of the findings from behavioral experiments. Some of the dysfunctional behavior that we encountered, motivated further histopathological and electrophysiological analysis of the hippocampus that pointed to a synaptopathy. In the end, we present the results of a single cell transcriptome screen that we conducted to identify differentially expressed genes that might be involved in the observed pathophysiology.

### Characteristic features and alterations of sleep patterns in *Pigv*^341E^ mice

The most prominent differences that we observed first between *Pigv*^341E^ and wild-type mice were reduced weight (Fig. 1B; SI *Appendix*, Fig. S1F) and hindlimb clasping behavior (Fig. 1C, D). Due to the intellectual disability and psychomotor delay that are the key clinical features of IGD, patients are impaired in their everyday lives. Therefore, we sought to determine which spontaneous behaviors our mouse model exhibited while living undisturbed in their home cage (singly or group-housed). Using HomeCageScan (HCS), we monitored singly-housed *Pigv*^341E^ mice for 23 h at two different time points (8 and 16 weeks). Among the 19 behaviors accurately detected by analysis of the HCS data, three behaviors were consistently altered in *Pigv*^341E^ mice at both time points. *Pigv*^341E^ mice hung less often to the top of the cage (total occurrences) and for shorter durations (total duration and duration per hour) than wild-type mice; groomed more often (total occurrences) and for longer durations (total duration and duration per hour); and slept less (total duration and duration per hour) (Fig. 2A, B; SI *Appendix*, Fig. S2A). At the earlier time point, *Pigv*^341E^ mice spent more time walking (duration per hour) during the dark phase of the day than wild-type mice (Fig. 2B, top graph). Furthermore, at both time points, hanging behavior was an important variable for differentiating genotypes along the dimensions of a principal component analysis (PCA) (SI *Appendix*, Fig. S4A–F).

**Figure 2.**
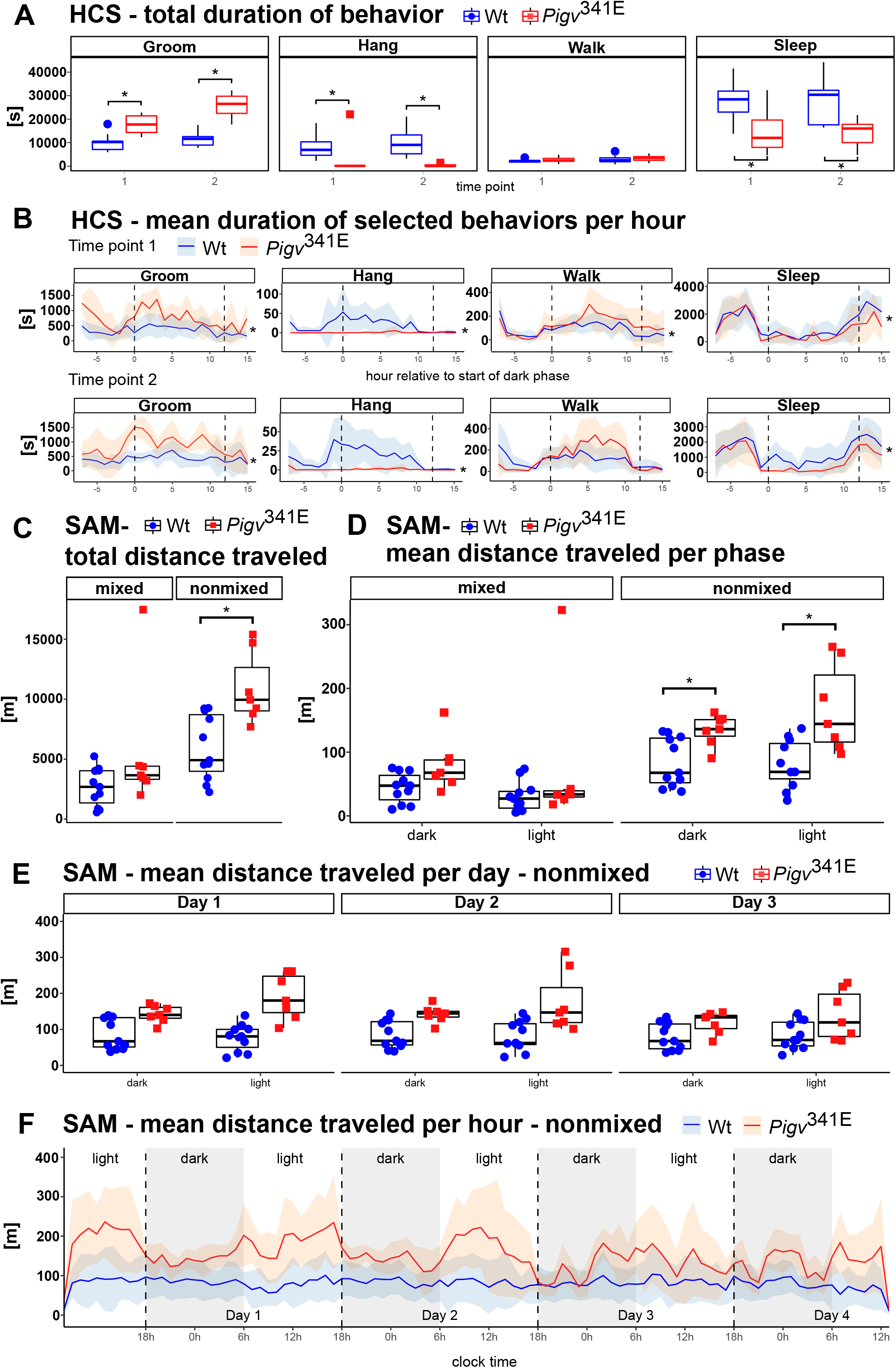
*Pigv*^341E^ mice exhibit a behavioral phenotype in HCS and an elevated activity level in the SAM (second approach). (A) Duration of selected behaviors from the homecage scan (HCS). *Pigv*^341E^ mice revealed elevated grooming and reduced hanging and sleeping behavior at both time points. (B) Duration of selected behaviors per hour, as determined by HCS. The dark phase took place between the two dashed vertical lines. *Pigv*^341E^ mice exhibited elevated grooming and hanging and reduced sleeping at both time points. *Pigv*^341E^ mice exhibited elevated walking at time point 1. (C) The social activity monitor (SAM) revealed an elevation in total distance traveled by *Pigv*^341E^ mice when animals were separated by genotype (non-mixed). When housed in mixed genotypes (mixed), total distance traveled did not significantly differ between genotypes. (D) *Pigv*^341E^ mice exhibited an increase in distance traveled per phase when separated by genotype (non-mixed). When housed in mixed genotype (mixed), total distance traveled per phase did not significantly differ between genotypes. (E, F) *Pigv*^341E^ mice traveled longer distances per day and per hour when separated by genotype (non-mixed). *Pigv*^341E^=homozygous for Pigv p.Ala341Glu; wt=wild type. Animals used for the HCS (wt: female n=8, male n=4; *Pigv*^341E^: female n=4, male n=2) were phenotyped at two different consecutive time points: tp1 at the age of 8 weeks, and tp2 at the age of 16 weeks. Animals used for the SAM: wt(female n=4, male n=6) *Pigv*^341E^(female n=2, male n=5). Data from the HCS (total duration) and SAM (total distance, mean distance traveled per phase and per day) were analyzed by Wilcoxon rank-sum test (non-parametric). Data from the HCS (duration per hour) and SAM (distance traveled per hour) were analyzed with a generalized linear mixed-effects model (glmm) using a Markov chain Monte Carlo (MCMC). *p < 0.05, **p < 0.01, ***p < 0.001.

We also used the social activity monitor (SAM) to assess spontaneous home-cage activity in *Pigv*^341E^ mice while living in a group setting. For this test, we implanted a RFID transponder into the mice and put the home cage with mixed genotypes on a grid box that could locate individual animals and their position in the cage at all times (continuous 24 h/day recording). Because the animals were undisturbed in their home cage, SAM analysis could be performed several times without the animals noticing. For the first two time points (9 and 17 weeks), SAM analysis revealed no difference between genotypes in total distance traveled over 14 days (SI *Appendix*, S4G), but an unexpected switch in diurnal/nocturnal activity for both genotypes was observed. The mice were more active during the light (normal sleeping time) than the dark phases of their days (SI *Appendix*, Fig. S5A, S5C; confirmed by Markov chain Monte Carlo generalized linear mixed-effects models [MCMCglmm)], pMCMC = 0.001). This observation supported the HCS data that *Pigv*^341E^ mice slept less, which is a feature of some patients with IGDs who suffer from sleep disturbances (7,20). We performed a second experiment in which we evaluated the spontaneous activity of the *Pigv*^341E^ and wildtype mice separately (non-mixed *vs*. mixed-genotype cages). When *Pigv*^341E^ and wild-type mice lived together in the same cage (mixed genotype), we reproduced our previous results: Mice of different genotypes exhibited no difference in total distance traveled over 4 days (Fig. 2C, left graph), per phase (Fig. 2D, left graph), per day (SI *Appendix*, Fig. 2C), and per hour (SI *Appendix*, Fig. S2D), but they still exhibited higher activity levels (larger variability) during the light *vs*. dark phase (SI *Appendix*, Fig. S2D, confirmed by Markov chain Monte Carlo generalized linear mixed-effects models [MCMCglmm], p = 0.001). However, in nonmixed housing conditions *Pigv*^341E^ mice walked greater distances over 4 days (Fig. 2C, right graph), per phase (Fig. 2D, right graph), per day, and per hour (Fig. 2E, F) than wild-type mice, and again, *Pigv*^341E^ mice were more active during the light than the dark cycle on days 1 and 2 (Fig. 2E).

### *Pigv*^341E^ mice exhibit motor dysfunction and alterations in sociability and social recognition

Extensive testing of various motor functions in *Pigv*^341E^ mice revealed a clear and elaborated dysfunctional motor phenotype. *Pigv*^341E^ mice had reduced balance and motor coordination, reflected by a reduced latency of falling off the rotarod (Fig. 3A). This was confirmed by an elevated latency of traversing an elevated beam (Fig. 3B). In the rope grip test, *Pigv*^341E^ mice exhibited an elevated latency of climbing on the rope, and had a lower hanging score than wild-type mice (Fig. 3C; SI *Appendix*, Fig. S6A right graph). Furthermore, *Pigv*^341E^ mice had reduced grip strength (Fig. 3D). Next, we evaluated the walking pattern of *Pigv*^341E^ mice using the footprint test (Fig. 3E). *Pigv*^341E^ mice had a larger distance between forepaw and hindpaw (S) in paw placement of the stride (Fig. 3F); this is remarkable because walking mice usually place their hind paws in the same positions as their forepaws (Fig. 3E, see wt).

**Figure 3.**
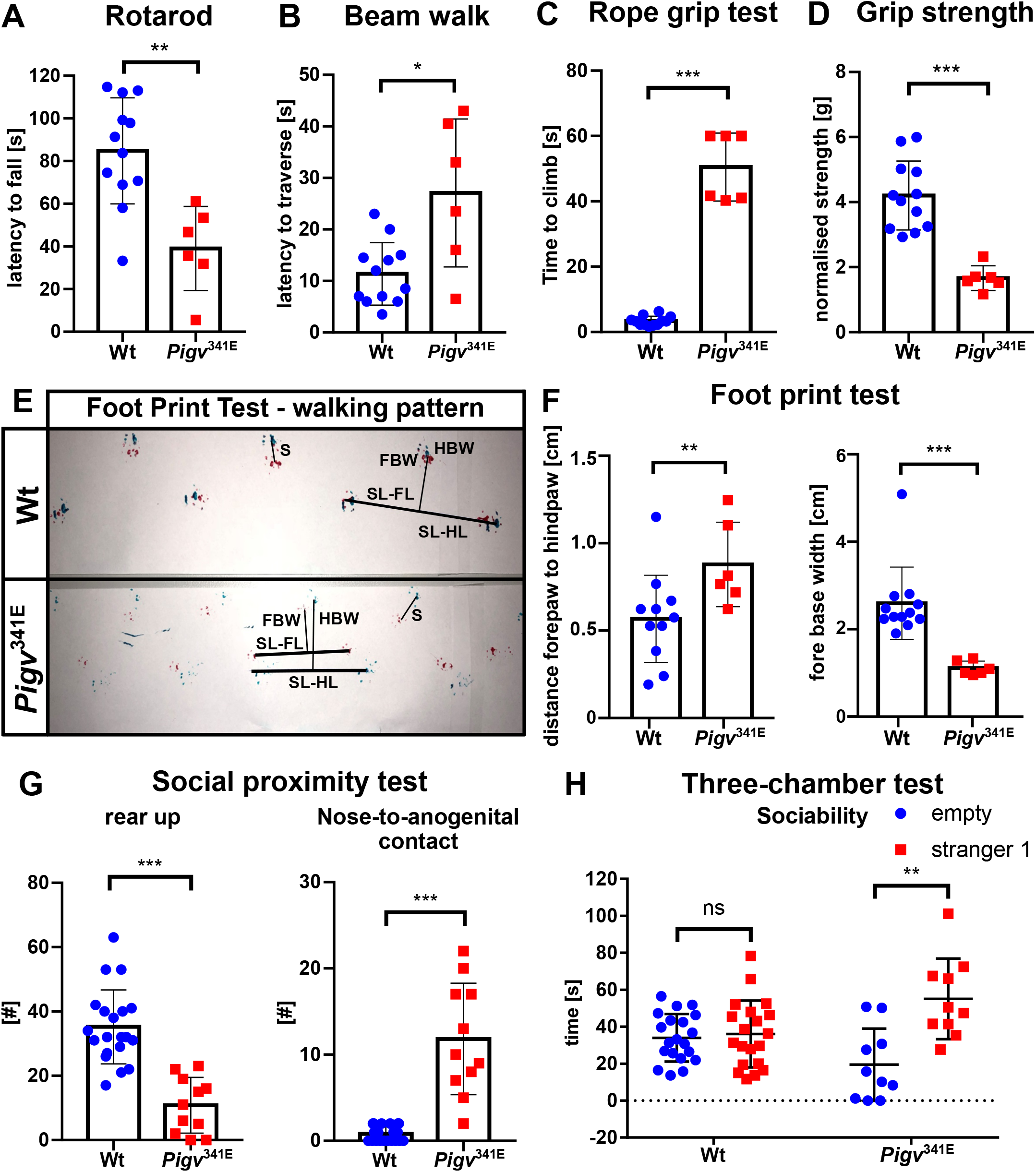
*Pigv*^341E^ mice exhibit motor dysfunctions. (A–B) *Pigv*^341E^ mice exhibited a reduction in motor coordination, reflected by a reduced latency of falling off the rotarod and elevated latency of traversal of the beam (diameter, 15 mm). (C) *Pigv*^341E^ mice exhibited a reduction in climbing performance, reflected by a decrease in the time spent climbing the rope. (D) *Pigv*^341E^ mice exhibited a reduced grip strength, as determined by grip strength meter. [g] was normalized to the weight of the animal. (E) Representative image of walking pattern between genotypes. SL–FL=stride length–forelimb, SL–HL=stride length–hindlimb, FBW=fore–base width, HBW=hind–base width, S=distance between forepaw and hindpaw. (F) *Pigv*^341E^ mice had an altered walking pattern, reflected by an increase in the distance between forepaw to hindpaw and a decrease in fore–base width. (G) *Pigv*^341E^ mice exhibited enhanced social approach behavior, reflected by a decrease in “rear up” behavior and an increase in the number of nose-to-anogenital contacts in the social proximity test. (H) The three-chamber test (first 5 min) indicated that *Pigv*^341E^ mice exhibited an enhanced social approach behavior relative to wild-type mice, reflected by spending more time with the stranger mouse than with the empty. *Pigv*^341E^=homozygous for Pigv p.Ala341Glu, wt=wild type. Animals used in the motor tests were 7 weeks old: wt(female n=8, male n=4) *Pigv*^341E^(female n= 4, male n=2). Animals used in the social proximity test: wt(female n=9, male n=11), *Pigv*^341E^(female n=4, male n=7). Animals used in the three-chamber test: wt(female n=9, male n=11), *Pigv*^341E^(female n=4, male n=6). The data from the motor tests and the social proximity test were analyzed by nonparametric t-test (Mann–Whitney). The data from the three-chamber test were analyzed by two-way analysis of variance (ANOVA) followed by Bonferroni’s multiple comparisons test. *p < 0.05, **p < 0.01, ***p < 0.001.

We evaluated social behavior in *Pigv*^341E^ mice in the social proximity and three-chamber test. In the social proximity test, *Pigv*^341E^ mice exhibited a reduced number of “rear up” behavior and an elevated number of nose-to-anogenital contacts with the stranger mouse (Fig. 3G). However, no differences in the number of nose-tip-to-nose-tip, nose-to-head-contact, “crawl over”, or “crawl under” behaviors were observed between genotypes (SI *Appendix*, Fig. S7A). In the three-chamber test, *Pigv*^341E^ mice spent more time with the stranger mouse than in the vicinity of the empty cage (Fig. 3H). Furthermore, the discrimination ratio (stranger *vs*. empty cage) was higher in *Pigv*^341E^ than in wild-type mice (SI *Appendix*, Fig. S7D, right graph). Nevertheless, in contrast to wild-type mice, *Pigv*^341E^ mice did not distinguish between the familiar (stranger 1) and unfamiliar mice (stranger 2) (SI *Appendix*, Fig. S7D, left graph).

### *Pigv*^341E^ mice exhibit cognitive deficits in spatial long-term memory and species-specific hippocampus-dependent functions

To characterize the cognitive and affective profile of *Pigv*^341E^ mice, we performed a battery of tests to assess aspects of spatial learning and memory, species-specific functions (Barnes maze, y-maze, marble burying, nest-building behavior), and spontaneous response to novel, open, and elevated or bright environments (open field, elevated plus maze, and dark/light-box). In the Barnes maze test, *Pigv*^341E^ mice were delayed in spatial learning, as indicated by an elevated latency to escape during days 1–3 (Fig. 4A). Despite this delay in spatial learning during days 1-3, *Pigv*^341E^ mice had learned the location of the escape box by day 4 and exhibited normal short-term spatial memory at day 5. However, *Pigv*^341E^ mice had impaired long-term spatial memory at day 12, reflected by an elevated latency to escape (Fig. 4A). Similar to the results that measured latency to escape, *Pigv*^341E^ mice had an elevated path length for days 1-4 and 12, but not day 5 (Fig. 4B). Furthermore, *Pigv*^341E^ mice spent less time in the target quadrant than wild-type mice at day 12, confirming deficits in long-term spatial memory (SI *Appendix*, Fig. S8B, right graph). In the y-maze test, similar spontaneous alternation behavior between genotypes suggested normal short-term spatial working memory in *Pigv*^341E^ mice (SI *Appendix*, Fig. S8A). In the species-specific and hippocampus-dependent tests of marble burying and nest construction (21), *Pigv*^341E^ mice buried fewer marbles and had lower-quality nests (Fig. 4C, D).

**Figure 4.**
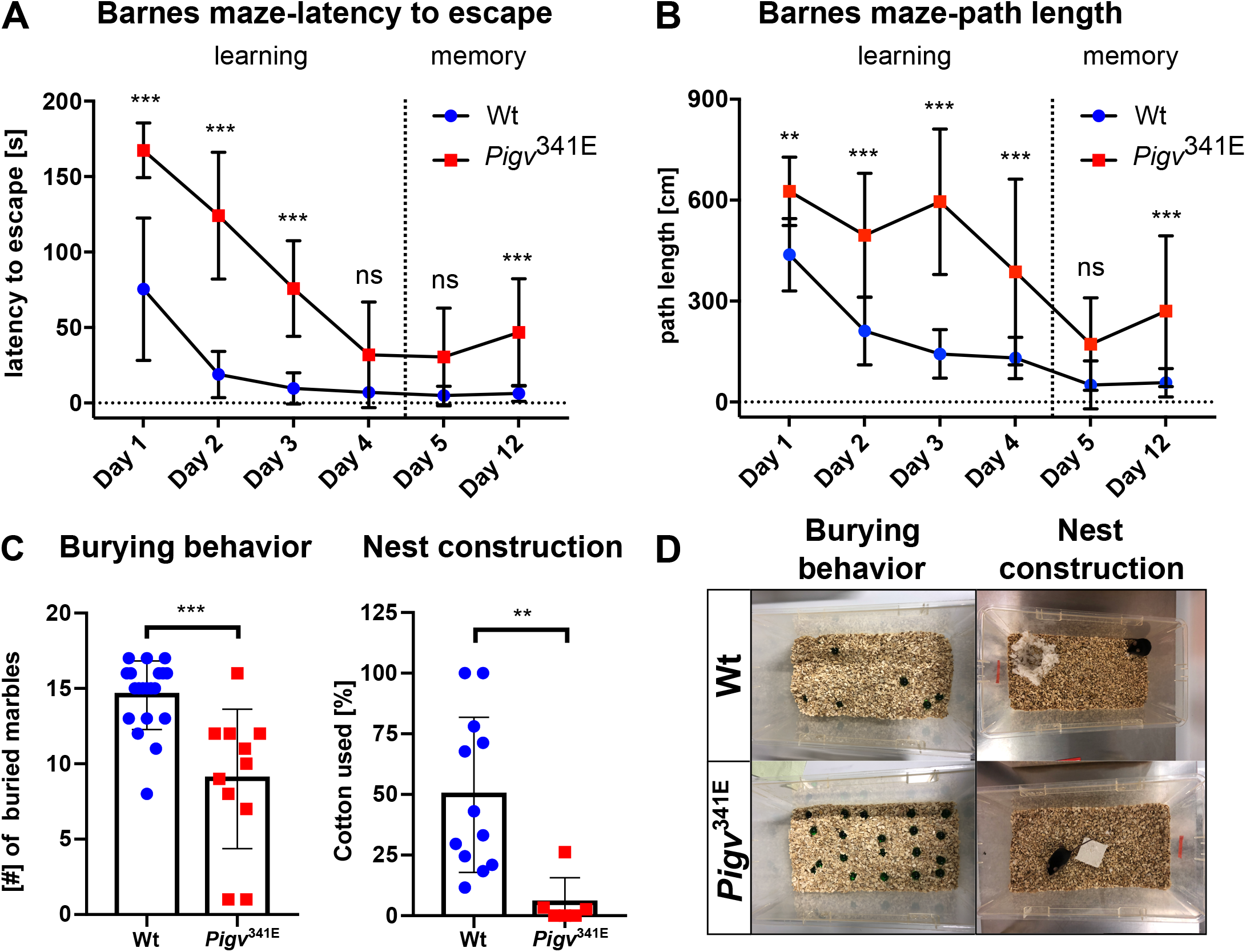
*Pigv*^341E^ mice exhibit cognitive deficits in learning and memory. (A–B) *Pigv*^341E^ mice exhibited cognitive deficits in learning and long-term memory, reflected by increases in latency to escape and path length on days 1–3 (learning) and day 12 (long-term memory in the Barnes Maze). (C) *Pigv*^341E^ mice exhibited a decrease in burying and nest construction behavior. (D) Representative image of burrowing behavior (left) and nest construction behavior (right) of wild-type and *Pigv*^341E^ mice. *Pigv*^341E^=homozygous for Pigv p.Ala341Glu; wt=wild type. Animals used in the Barnes maze: wt(female n=8, male n=11), *Pigv*^341E^(female n=4, male n=6). Animals used in the marble-burying test: wt(female n=9, male n=11), *Pigv*^341E^(female n=4, male n=7). Animals used in the nest construction test were 7 weeks old: wt(female n=8, male n=4), *Pigv*^341E^(female n=4, male n=2). The data from the nest construction and marble-burying tests were analyzed with a non-parametric t-test (Mann–Whitney). The data from the Barnes maze (latency to escape, path length) were analyzed by two-way analysis of variance (ANOVA) followed by Bonferroni’s multiple comparisons test. *p < 0.05 **p < 0.01, ***p < 0.001.

*Pigv*^341E^ mice behaved similarly to wild-type mice in the dark/light-box and elevated plus maze (SI *Appendix*, Fig. S9A, B). In the open field test, *Pigv*^341E^ mice spent less time in the center and more time in the periphery than wild-type mice (SI *Appendix*, Fig. S9C). Moreover, the path length and number of visits to the center were reduced in *Pigv*^341E^ mice (SI *Appendix*, Fig. S9D, E).

### *Pigv*^341E^ mice exhibit defects in synaptic transmission

Because *Pigv*^341E^ mice exhibited cognitive impairments in spatial learning and memory, we hypothesized a hippocampal defect, and this idea was supported by the reduced burrowing and nest-building behavior (hippocampus-dependent) in *Pigv*^341E^ mice. Hence, we decided to analyze the hippocampus in more detail. Nissl staining revealed no morphological abnormalities in the hippocampus of *Pigv*^341E^ mice (SI *Appendix*, Fig. S10C). Because many GPI-linked proteins play crucial roles in synapse formation and plasticity, causing deficits that can be detected in cell culture (22), we performed immunostaining to visualize synaptophysin, a presynaptic marker. We observed a decreased immunoreactivity for synaptophysin in *Cornu Ammonis 1–Stratum Radiatum* (CA1–SR) of *Pigv*^341E^ mice (Fig. 5A). By contrast, we observed no significant differences between genotypes in *Cornu Ammonis* 3–*Stratum Radiatum* (CA3–SR) or *Cornu Ammonis* 1–molecular layer of dentate gyrus (CA1–ML) (SI *Appendix*, Fig. S10A, B). To determine whether the behavioral abnormalities were accompanied by deficits in synaptic transmission, we conducted electrophysiology recordings in the CA1–SR region, where synaptophysin levels were reduced in *Pigv*^341E^ mice. We tested input–output functions and observed lower amplitudes of excitatory post-synaptic potential (EPSPs) at different stimulation intensities in *Pigv*^341E^ mice (Fig. 5B). In paired pulse facilitation (PPF), *Pigv*^341E^ mice exhibited an elevated paired pulse ratio (PPR) (Fig. 5C). Moreover, post-tetanic potentiation (PTP) exhibited elevated facilitation in *Pigv*^341E^ mice (Fig. 5D).

**Figure 5.**
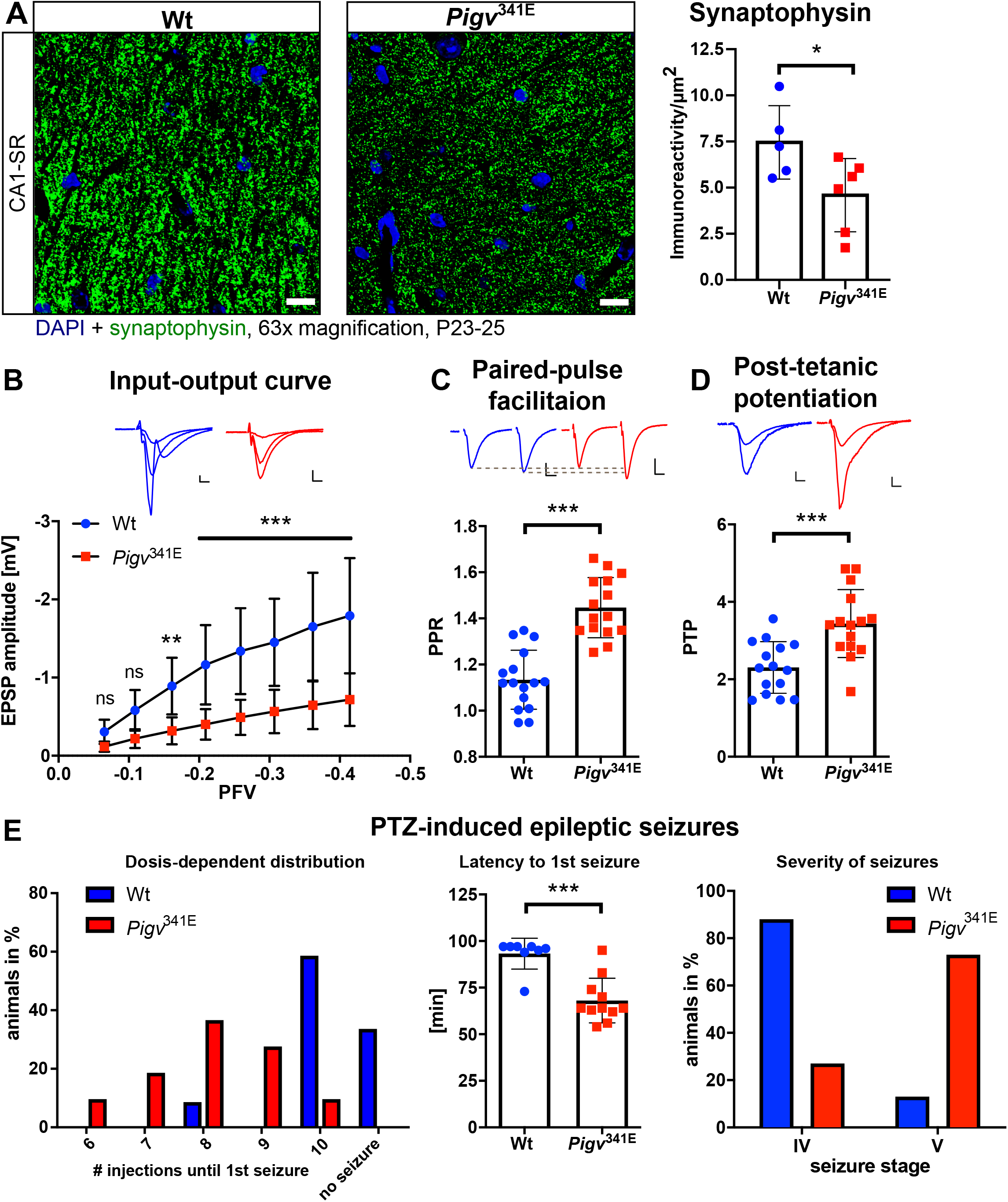
*Pigv*^341E^ exhibit a hippocampal synaptic defect. (A) Representative images of immunofluorescence staining show less synaptophysin immunoreactivity (green) in the *Cornu Ammonis 1–Stratum Radiatum* (CA1–SR) of *Pigv*^341E^ mice. 4’,6-diamidino-2-phenylindole (DAPI) staining is shown in blue. Scale: 10 μm. Synaptophysin immunoreactivity per μm^2^, quantified using FIJI ImageJ, was significantly reduced in CA1–SR in *Pigv*^341E^ mice (right). (B) Input–output: excitatory post-synaptic potential (EPSP) amplitude was reduced in *Pigv*^341E^ mice at different presynaptic fiber volleys (PFV). Scale: L indicates 0.2 mV/5 ms. (C) Paired pulse facilitation (PPF) revealed an elevated paired pulse ratio (PPR) in *Pigv*^341E^ mice. Scale: L indicates 0.2 mV/10 ms. (D) Post-tetanic potentiation (PTP) was elevated in *Pigv*^341E^ mice. Scale: L indicates 0.2 mV/5 ms. (E) In the PTZ kindling model *Pigv*^341E^ mice had a significantly different seizure threshold compared to that of wildtype control littermates (left graph). In *Pigv*^341E^ mice the latency to the first convulsive seizure was significantly reduced (middle graph). Furthermore, the seizure phenotype appears more severe (right graph). *Pigv*^341E^=homozygous for Pigv p.Ala341Glu; wt=wild type. Animals used for the synaptophysin-staining were 3 weeks old: wt(female n=3, male n=2) *Pigv*^341E^(female n=2, male n=4). Animals used for the electrophysiology recordings: wt(female n=2, male n=2) *Pigv*^341E^(female n=2, male n=2). Animals used for the PTZ model: Wt(female n=3, male n=9) *Pigv*^341E^(female n=5, male n=6). The data from the synaptophysin immunofluorescence staining and electrophysiology recordings (PPF, PTP) were analyzed by parametric Student’s t-test. Furthermore, the data from input-output curve were analyzed by two-way analysis of variance (ANOVA) followed vy Bonferroni’s multiple comparisons test. In the PTZ model, group comparisons (dosis-dependent distribution, severity of seizures) were analyzed by Chi-squared test and latency to first seizure by unpaired student t-test with Welch’s correction. *p < 0.05 **p < 0.01, ***p < 0.001.

### Increased susceptibility of *Pigv*^341E^ mice to chemically induced acute seizures

Considering the altered electrophysiological properties of in *Pigv*^341E^ mice, we examined for differences in the seizure threshold between *Pigv*^341E^ and corresponding littermate wild-type mice in an acute epilepsy model, the pentylenetetrazole (PTZ)-induced kindling model (23). To this end, mice were repetitively exposed to PTZ (10 mg/kg, i.p.) every 10 min until the occurrence of a first focal to bilateral tonic-clonic seizure occurred. Intriguingly, *Pigv*^341E^ mice exhibited a significantly lower seizure threshold manifesting as a reduced latency to first seizure than wild-type mice which exhibited convulsive seizures after PTZ exposure. Wildtype animals manifested generalized seizures after 93.3 min; whereas in *Pigv*^341E^ mice, the first seizure manifested already after 63.3 min (Fig. 5E, middle graph). Furthermore, the observation that four wild-type animals did not exhibit any seizure after 10 injections, whereas all *Pigv*^341E^ mice did, underlines the higher susceptibility of *Pigv*^341E^ mice for seizure induction via PTZ (Chi-Square= 105.3, df=4, p<0.0001) (Fig. 5E, left graph). In addition, *Pigv*^341E^ mice exhibited more severe seizures than wild-type mice (Chi-Square=74.21, df=1, p<0.0001) (Fig. 5E, right graph).

### Shift in relative cell count of hippocampal cellular subgroups in *Pigv*^341E^ mice

The synaptic defect in the CA1–SR region of *Pigv*^341E^ mice could have been responsible for the observed impairments in spatial learning and memory. To identify the cell types most affected by GPIBD, we performed single-cell RNA sequencing on freshly isolated hippocampal cells (24) after cognitive behavioral tests. Based on the gene expression profiles of 8,800 single-cells from *Pigv*^341E^ mice and 7,100 cells from wild-type animals, we defined 17 cellular subgroups (Fig. 6A). Cells from both *Pigv*^341E^ and wild-type mice were present in all subgroups, but the distributions differed between genotypes (*χ*^2^ = 306.49, *df* = 16, *p* < 2.2 * 10^−16^). While the fractions of granule cells, oligodendrocytes, and a microglia subpopulation (microglia 3) were reduced in pooled samples from *Pigv*^341E^ mice, the proportions of subicular neurons (neurons subiculum 1), GABAergic (inhibitory) interneurons, and fibroblast-like cells were higher than in the pooled wild-type samples (Fig. 6B).

**Figure 6.**
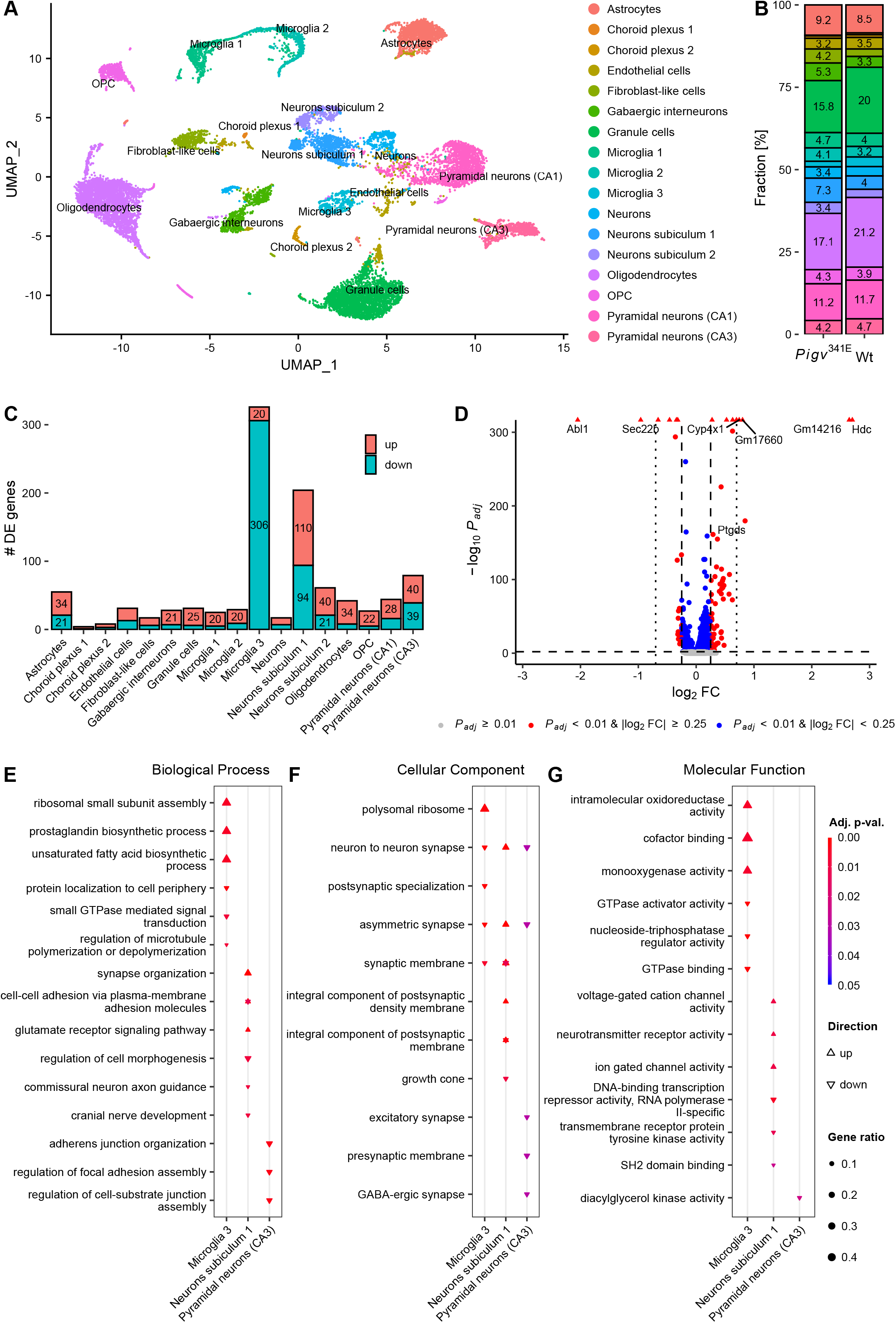
Differences between *Pigv*^341E^ and wild-type mice in the transcriptional landscape of single hippocampus cells. (A) Hippocampus cells from both *Pigv*^341E^ and wild-type mice clustered into 17 cellular subgroups (B), with a skewed distribution of cells across subgroups in the *Pigv*^341E^ sample relative to wild type. Percentages below 3% are not shown for clarity. (C) Differential expression testing between *Pigv*^341E^ and wild-type cells within each cellular subgroup yielded differentially expressed genes with absolute log_2_(fold change) ≥ 0.25 and adjusted p-value ≤ 0.01. The most extensive changes in gene expression were observed for the third subgroup of microglia cells and the first subgroup of subicular neurons. Gene counts are not shown for sets with < 20 differentially expressed genes. (D) Global comparison between all *Pigv*^341E^ and wild-type cells highlights the hippocampus-wide effect on the genes *Abl1, Hdc, Cyp4x1*, and *Gm14216*, for which significant changes in expression were detected within each cellular subgroup. The dashed horizontal line is located at an adjusted p-value of 0.01, the dashed vertical lines at an absolute log_2_(fold change) of 0.25, and the dotted vertical lines at an absolute log_2_(fold change) of 0.7. For genes with an absolute log_2_(fold change) > 0.7, gene symbols are shown. Within the subgroups of microglia 3, neurons subiculum 1 and pyramidal neurons (CA3), GO enrichment analysis revealed a number of over-represented Biological Process (E), Cellular Component (F), and Molecular Function terms (G) in the sets of genes differentially expressed between *Pigv*^341E^ and wild-type cells within the cellular subgroups. For each set of genes that were upregulated or downregulated between genotypes, up to three terms with the lowest adjusted p-value after removal of redundant GO terms are shown. The gene ratio, i.e., the number of genes annotated with a term divided by the respective number of differentially expressed genes, is encoded by point size. Cellular subgroups with no significantly enriched terms were omitted. GO terms enriched in differentially expressed genes with elevated or reduced expression within a subgroup are contracted into a single starshaped symbol. DE genes, differentially expressed genes; FC, fold change; GO, Gene Ontology; P_adj_, adjusted p-value. Pooled samples [wt(male=4), *Pigv*^341E^(male=4)]. n=1 per genotype.

### *Pigv*^341E^ hippocampal cells exhibit a de-regulation in gene expression related to synapse organization and signaling transduction

Expression analysis of single-cell RNA sequencing data revealed multiple genes that were differentially expressed between *Pigv*^341E^ and wild-type cells within each cellular subgroup (Fig. 6C). In the mutant mice, non-receptor tyrosine kinase *Abl1* was downregulated, whereas histidine decarboxylase *Hdc*, cytochrome P450 member *Cyp4x1*, and predicted lncRNA *Gm14216* were upregulated, both within and across all cellular subgroups (Fig. 6D; SI *Appendix*, Fig. S16–18). The most extensive change in gene expression within a cellular subgroup was observed in the first subgroup of subicular neurons and the third subgroup of microglia (Fig. 6C; SI *Appendix*, Fig. S16–17). We also performed Gene Ontology (GO) analysis of differentially expressed genes between genotypes, including terms in three categories: Biological Process, Cellular Component, and Molecular Function (Fig. 6E–G). Among the differentially expressed genes in subicular neurons, the Biological Process term “regulation of synapse organization” was enriched in genes upregulated in *Pigv*^341E^ cells, whereas the Biological Process terms “cell morphogenesis” and “commissural neuron axon guidance” were enriched in downregulated genes (Fig. 6E). Furthermore, the Biological Process term “cell–cell adhesion via plasma-membrane adhesion molecules” was enriched in genes with elevated and reduced expression (Fig. 6E).

Single-cell RNA sequencing revealed three hippocampal microglia populations in both genotypes (Fig. 6A, B, microglia 1–3). All three subpopulations expressed the marker genes *Csf1r* and *C1qa* (SI *Appendix*, Fig. S15). Among the most significant differentially expressed genes between the three microglia subgroups and all remaining cells in both genotypes, “cell activation”, “migration”, “phagocytosis”, and “immune responses” were among the top 10 GO Biological Process terms in microglia 1 and 2 (SI *Appendix*, Dataset S2). By contrast, the top 10 GO Biological Process terms in microglia 3 cells were “ribosome”, “ribonucleoprotein complex biogenesis”, and “cytoplasmic translation” (SI *Appendix*, Dataset S2). Hence, we considered microglia 1 and 2 cells as potentially more phagocytic and migratory than microglia 3 cells. We identified 326 genes that were differentially expressed between genotypes in microglia 3 cells. Remarkably, 306 of these 326 genes were downregulated in microglia 3 cells of *Pigv*^341E^ mutants (Fig. 6C). GO analysis revealed that the downregulated genes were enriched for the Biological Process terms “small GTPase-mediated signal transduction” and “regulation of microtubule cytoskeleton polymerization and depolymerization” (Fig. 6E). In addition, we identified 20 genes differentially expressed between genotypes that were upregulated in *Pigv*^341E^ microglia 3 cells. Among these were *Rp138, Rps21*, and *Rps28*, which encode ribosomal proteins.

CA3 pyramidal neurons project their axons to the CA1 region, where we observed dysfunction in synaptic transmission in *Pigv*^341E^ mice. Therefore, we were particularly interested in the cellular subgroup of CA3 pyramidal neurons, in which we identified 79 genes that were differentially expressed between genotypes (Fig. 6C). GO analysis revealed that downregulated genes were enriched for the terms “adherens junction organization”, “regulation of cell-substrate junction assembly”, and “focal adhesion assembly” in *Pigv*^341E^ CA3 pyramidal neurons (Fig. 6E). Interestingly, we also observed enrichment of genes with reduced expression related to “neuron–neuron synaptic transmission”, “regulation of synaptic vesicle exocytosis”, and “synaptic vesicle transport” (SI *Appendix*, Dataset S3).

## Discussion

*Pigv*^341E^ is the first mouse model for GPI-anchor deficiency with a hypomorphic mutation that is viable after weaning. The mice exhibited significant alterations in behavior that reflect key aspects of patients with IGD.

In these mice, we observed a severe motor phenotype that included deficits in motor coordination, grip or muscle strength, climbing, and hanging behavior (in HCS); alterations in walking pattern; and hindlimb clasping. Behavioral traits such as altered walking pattern, hindlimb clasping, and motor coordination deficits are usually observed in mouse models with ataxia-like behavior and cerebellar dysfunction (25,26). In agreement with these findings, ataxia has been reported in some IGD patients, and an ataxia-like behavior was observed in a conditional *Piga* knockout mouse model (10,19). Lukacs et al. analyzed the microscopic anatomy of cerebellum sections from their conditional *Piga* knockout mouse model and observed mild deficits in Purkinje cell arborization (10). However, in histologic analysis with various stainings, the cerebellum of our *Pigv*^341E^ mutants did not exhibit any abnormalities (SI *Appendix*, Fig. S11A–F), and the overall folial pattern appeared to be unchanged. In particular, calbindin staining exhibited no differences in Purkinje cell dendritic arborization between genotypes (SI *Appendix*, Fig. S11E–F). Therefore, we hypothesize that deficits in dendritic arborization in this neuronal cell type, as reported by Lukacs et al., require a more severe GPIBD than that induced by the hypomorphic mutation c.1022C>A in *Pigv*. This is consistent with the longer lifespan of our mouse model, which allowed us to analyze the associated cognitive deficits. In addition to the cerebellum, we focused on the hippocampus, where we performed histology and electrophysiology to achieve a deeper understanding of the memory and species-specific deficits. Although we did not observe significant morphological changes in the hippocampus, the input–output curve, PPR, and PTP were significantly altered in *Pigv*^341E^ mice, indicating that electrophysiology is a sensitive functional assay for mouse models with mild GPIBD.

Because *Pigv*^341E^ mice exhibited increased self-grooming, which is a repetitive, highly stereotyped pattern that is associated with autistic-like behavior in rodents (27), we suspected abnormalities in social behavior as well. Autistic features have been reported in a subgroup of patients with IGD due to pathogenic mutations in *PGAP3* (7). By contrast, patients with *PIGV* deficiency are keen to interact socially despite their severe speech impairments. Interestingly, in *Pigv*^341E^ mice we observed enhanced social approach behavior, reflected by an elevated number of nose-to-anogenital contacts and reduced “rear up” behavior in the social proximity test. The reduction in “rear up” behavior in *Pigv*^341E^ mice suggested reduced social avoidance.

The enhanced social approach behavior was confirmed in the three-chamber test: relative to wild-type controls, *Pigv*^341E^ mice spent more time with the stranger mouse than with the empty cage. Comparable performance between genotypes in the buried food test ruled out compromised olfaction, a potential confounder in social behavior tests (SI *Appendix*, Fig S7C). Taking into account the enhanced social approach behavior in *Pigv*^341E^ mice, the positive social abilities of patients with *PIGV* deficiency seem to be characteristic of these individuals and should be considered during diagnosis. However, it remains unknown to what extent IGD patients who are affected in genes other than *PIGV* exhibit positive social abilities. In addition, because we did not observe social behavior characteristic of autism (28) in *Pigv*^341E^ mice, and autistic features are seen only in a subgroup of patients with IGD, autism should not be considered as a specific feature of IGD.

*Pigv*^341E^ mice exhibited a deficit in spatial long-term memory in the Barnes maze, correct short-term spatial memory, and short-term working memory (y-maze test). Furthermore, *Pigv*^341E^ mice exhibited a delay in spatial learning relative to wild-type mice in the Barnes maze (day 1–3). In the Barnes maze, both latency to escape and path length were elevated in *Pigv*^341E^ mice; therefore, we excluded the motor phenotype as a confounder. However, even though path length and latency to escape were elevated in *Pigv*^341E^ mice, the number of visits to the wrong holes was not significantly altered on days 2 through 12 (SI *Appendix*, Fig. S8C). Analysis of the search strategy revealed that *Pigv*^341E^ mice were targeting the correct hole without any errors (number of wrong holes) less often than wildtype mice (SI *Appendix*, Fig. S8D, direct strategy) (Wt: 15.8 %, *Pigv*^341E^: 3.3%). Moreover, we quantified two further search strategies: random choice of a hole without any order, and a serial strategy that tests holes one after another in close proximity. *Pigv*^341E^ mice used more often the random strategy (Wt: 9.7 %, *Pigv*^341E^: 56.7 %), whereas wild-type mice used more often the serial strategy to find the correct hole (Wt: 25.4 %, *Pigv*^341E^: 20.0 %) (SI *Appendix*, Fig. S8D). In contrast to random guessing, the serial strategy has the advantage that it minimizes the path length and by that means will result in a quicker escape. However, the serial strategy results also in a higher number of errors. This potentially explains why the latency to escape was significantly elevated in *Pigv*^341E^ mice, whereas the number of wrong holes did not significantly differ between genotypes.

Furthermore, we observed no difference between genotypes in affective related behavior (dark/light-box, elevated plus maze) except in the open field test, in which *Pigv*^341E^ mice spent more time in the periphery than in the center. However, this observation could also represent a confounder due to the motor phenotype, as the number of entries to the center and the path length were also reduced in *Pigv*^341E^ mice.

Interestingly, electrophysiology recordings revealed reduced synaptic transmission at CA1–SR in *Pigv*^341E^ mice, consistent with the observed impairment in long-term spatial memory and hippocampus-dependent species-specific behaviors (marble-burying and nest construction test). While PTP and PPR were elevated in *Pigv*^341E^ mice, the input–output curve was reduced, indicating a decrease in synaptic release probability. Due to the increase in PTP and PPR, and the reduced immunoreactivity of synaptophysin, a presynaptic vesicle protein, we hypothesize that the pool of readily releasable vesicles in the pre-synapse is reduced, resulting in a damped input–output curve in the post-synapse. Notably in this regard, GO analysis of single-cell RNA sequencing data revealed that genes which were significantly downregulated in *Pigv*^341E^-CA3 pyramidal neurons were enriched for GO Biological Process terms associated with “synaptic transmission” and “vesicle transport”. The impaired synaptic transmission in the hippocampi of *Pigv*^341E^ mice was reflected in their lower threshold to the excitotoxin-induced epileptic events and aggravated seizures than wild-type mice.

Single-cell RNA sequencing data revealed the most prominent differences in gene expression in a subgroup of subicular neurons and microglia. In subicular neurons 1, upregulated genes were associated with Biological Process terms such as “synapse organization”. Because we observed a synaptic defect in CA1–SR, as revealed by immunohistochemistry and as further supported by electrophysiological recordings, we hypothesized a synaptic defect in the subiculum as well. The subiculum, an area of the hippocampus that is important for memory retrieval, is linked through microcircuits with the CA1 (29). Lederberger et al. described two distinct circuits for memory acquisition and retrieval: memory acquisition involves the CA1 and medial-entorhinal-cortex, whereas memory retrieval involves the CA1, the medial-entorhinal-cortex, and the subiculum. Future studies should seek to determine whether memory acquisition, memory retrieval, or even both conditions are affected in *Pigv*^341E^ mice.

Strikingly, 306 genes were downregulated in *Pigv*^341E^ microglia 3 cells. Therefore, microglia might play more important roles in GPI-anchor deficiency than previously thought. These genes were enriched in GO Biological Process terms “protein localization to cell periphery”, “small GTPase-mediated signal transduction”, and “regulation of microtubule polymerization or de-polymerization”. Small GTPases are important mediators of the cytoskeleton (30). Hence, we hypothesized that a GPI-anchor defect leads to downregulation of small GTPase-mediated pathways, which has further consequences for cytoskeleton organization in this microglia subtype. In this regard, GPI-anchored ephrin A proteins could play an important role, as EphrinA1 regulates small GTPase (Rho)-dependent cytoskeleton rearrangement through Src/focal adhesion kinases (31).

Up to 0.5% of the eukaryotic proteins are GPI-linked, with a broad range of functions including cell–cell adhesion, signal transduction, and antigen presentation (32). Therefore, it is surprising that the pathophysiology of acquired GPI-anchor deficiency PNH can be explained by the reduced expression of only two substrates, CD55 and CD59, which reduces the protection of cells against membrane attack complex (MAC), and can also be effectively treated by eculizumab, which inhibits complement activation (33). Likewise, analysis of mouse models of congenital forms of GPI-anchor deficiencies has aimed at identifying other lead targets. McKean et al. suggested a pivotal role of Cripto/TGFβ signaling in the development of holoprosencephaly (14), whereas Lukacs et al. discussed the role of GPI-anchored Folr1 in neural crest cells and the cranial neuroepithelium, and argued that compromise of Folr1 could be linked to the facial gestalt (13).

Interestingly, a considerable number of GPI-linked proteins are involved specifically in synapse formation and plasticity (1). Because *Pigv*^341E^ mice exhibit a hippocampal synaptic defect, this subset of GPI-linked proteins, including GPI-linked EphrinA, may play pivotal roles in the development of the disease as well. Single-cell RNA sequencing analysis revealed that *Abl1*, which interacts on the protein level with several EphrinA receptors (SI *Appendix*, Fig. S6F) (34), was not only downregulated in *Pigv*^341E^ mice across all cellular subgroups, but also within each cellular subgroup. Our hypothesis is that the GPI-anchor defect in the hippocampus is especially critical for EphrinA signaling, and that defective GPI anchoring of EphrinA in turn reduces hippocampal EphrinA receptor and Abl1 activity. Notably in this regard, axon repulsion is EphrinA-dependent and mediated through the Abl kinase family (35)..

Along with *Abl1, Hdc* and *Ptgds* were also dysregulated in *Pigv*^341E^ mice, with elevated expression across all cellular subgroups. *Hdc* encodes a histidine decarboxylase that catalyzes the conversion from histidine to histamine, an important neurotransmitter regulating circadian rhythm (36). In rodents, histamine levels are elevated during the dark phase to induce wakefulness and are reduced during the light phase to induce sleep (37). Furthermore, intracerebroventricular application of histamine triggers characteristic signs of wakefulness, such as elevated grooming and exploration behavior, which were also observed in *Pigv*^341E^ mice (HCS, SAM) (37). Furthermore, during home-cage activity monitoring (group-housed) and the HCS (individually housed), *Pigv*^341E^ mice were more active during the light cycle and slept for shorter durations. Consequently, higher expression of *Hdc* may lead to higher production of histamine, thereby disturbing circadian rhythm and causing classical signs of wakefulness in *Pigv*^341E^ mice during the light phase. However, it remains unknown how misregulation of *Hdc* is associated with GPI-anchor deficiency. Interestingly, in the conditional *Piga* knockout mouse model, bulk RNA sequencing of the cerebellum revealed an enrichment of de-regulated genes associated with the circadian rhythm (10). Moreover, *Ptgds* encodes prostaglandin D2 synthase, which converts prostaglandin H2 (PGH2) into prostaglandin D2 (PGD2), which in turn induces sleep (38). PGD2 levels fluctuate with the circadian rhythm and are elevated in the cerebrospinal fluid when rats are sleep-deprived (39). Therefore, upregulation of *Ptgds* expression could be an indicator of sleep deprivation in *Pigv*^341E^ mice.. Because *Pigv*^341E^ mice exhibit fewer resting phases during their classical inactive (light) phase, a careful analysis of sleep pattern in IGD patients is indicated. To date, sleep disturbances have mainly been reported in patients with *PGAP3* deficiency (19).

In summary, we have performed a deep phenotypic characterization of a mouse model that mirrors the symptoms of human patients with IGD. In addition, we detected a hippocampal synaptic defect that may impair spatial long-term memory and important species-specific behaviors for the survival of the animal. We hope that our model, as well as our phenotyping approach, will be useful in future studies aimed at a detailed elucidation of the pathomechanism of IGD and the response to therapeutic interventions.

## Materials and Methods

For full methods, see supplementary Material and Methods (SI *Appendix*).

### Animals

*Pigv*^341E^ mice were generated by diploid or tetraploid aggregation (40) (SI *Appendix*, Fig. S1A) and maintained by crossing with C57Bl.6/J mice. Mice were genotyped by PCR using the primers m*Pigv*Ex4_fw and m*Pigv*Ex4_rv. PCR amplicons were digested with *BcuI* (Thermo Fisher Scientific) and subjected to agarose gel electrophoresis (SI *Appendix*, Fig. S1D). All animals were handled according to government regulations as approved by local authorities (LaGeSo Berlin and LANUV Recklinghausen). In addition, all experiments were carried out following the 3R guidelines for animal welfare. Mice were housed in groups with mixed genotypes in single ventilated cages with an enriched environment. The mice were housed in a pathogen-free animal facility with a 12 h dark/light cycle, and had food and water *ad libitum* unless otherwise indicated. Mice used for experiments were 8 weeks to 6 months old unless otherwise indicated. *Pigv*^341E^ and wild-type mice used in a given experiment were the same age. To avoid bias effects, littermates were assigned equally to both experimental groups, according to their genotype and sex. The experimenter was blinded except during behavioral testing, as *Pigv*^341E^ mice were physically smaller. Moreover, experiments were randomized with respect to mouse genotype.

### Statistical analysis

For all experiments, at least four animals per genotype were used, except for the weight curve, for which at least three animals per genotype were used. One animal was defined as one biological replicate and represented one data point, except for electrophysiology recordings (see section: Schaffer collateral recordings). Means and standard deviations were calculated for each genotype group unless otherwise indicated. Data were statistically analyzed with GraphPad Prism (Version 7) or R (41), and results are expressed as means ± standard deviations. Statistical tests were performed for each experiment as indicated in SI *Appendix*, Dataset S1. Results with p-value < 0.05 were considered significant unless otherwise indicated.

## Supporting information

SI-Supplementary Dataset 1-statistical analysis

SI-Supplementary Dataset 2-GO_cellsubset

SI-Supplementary Dataset 3-GO_genotype_genotypeByCellsubset

SI-Supplementary Dataset 4-DEG_cellsubset

SI-Supplementary Dataset 5-DEG_genotype_genotypeByCellsubset

SI-Supplementary Dataset 6-motor tests

SI-Supplementary Dataset 7-social tests

SI-Supplementary Dataset 8-Barnes maze

SI-Supplementary Dataset 9-cognitive, species-specific tests

SI-Supplementary Dataset 10-FACS, weight curve, hindlimb-clasping

SI-Supplementary Dataset 11-Synaptophysin-Immunoreactivity

SI-Supplementary Dataset 12-electrophysiology_PTZexperiments

SI-Supplementary Dataset 13-SocialActivityMonitor_1stApproach

SI-Supplementary Dataset 14-SocialActivityMonitor_2ndApproach

SI-Supplementary Dataset 15-HomeCageScan

SI-Supplementary Dataset 16-affective behavior

## Data availability

The single-cell RNA sequencing data are freely available from the Gene Expression Omnibus (GEO) repository under accession number GSE147722. Other data are available from the corresponding author on reasonable request.

## Acknowledgments

We thank the Core Unit for Bioinformatics Data Analysis (CUBA) from the University Hospital of Bonn for their support. We thank Andrea Christ from the Anatomisches Institut, Anatomie & Zellbiologie, Medical Faculty of the University of Bonn, for excellent technical support. Furthermore, we thank the Animal Outcome Core Facility (AOCF) of the NeuroCure center and Charité-Universitätsmedizin, as well as the animal facility, transgenic core, and the sequencing core facility from the Max Planck Institute for Molecular Genetics in Berlin for their support.

## References

1. Um JW, Ko J. Neural Glycosylphosphatidylinositol-Anchored Proteins in Synaptic Specification. Trends Cell Biol [Internet]. 2017;27(12):931–45. Available at: http://dx.doi.org/10.1016/j.tcb.2017.06.007

2. Paulick MG, Bertozzi CR. The glycosylphosphatidylinositol anchor: A complex membrane-anchoring structure for proteins. Biochemistry. 2008;47(27):6991–7000.

3. Takeda J, Miyata T, Kawagoe K, Iida Y, Endo Y, Fujita T, et al. Deficiency of the GPI anchor caused by a somatic mutation of the PIG-A gene in paroxysmal nocturnal hemoglobinuria. Cell. 1993;73(4):703–11.

4. Bellai-Dussault K, Nguyen TTM, Baratang N V., Jimenez-Cruz DA, Campeau PM. Clinical variability in inherited glycosylphosphatidylinositol deficiency disorders. Clin Genet. 2019;95(1):112–21.

5. Murakami Y, Nguyen TTM, Baratang N, Raju PK, Knaus A, Ellard S, et al. Mutations in PIGB Cause an Inherited GPI Biosynthesis Defect with an Axonal Neuropathy and Metabolic Abnormality in Severe Cases. Am J Hum Genet. 2019;105(2):384–94.

6. Knaus A, Kortüm F, Kleefstra T, Stray-Pedersen A, Ðukić D, Murakami Y, et al. Mutations in PIGU Impair the Function of the GPI Transamidase Complex, Causing Severe Intellectual Disability, Epilepsy, and Brain Anomalies. Am J Hum Genet. 2019;105(2):395–402.

7. Knaus A, Awaya T, Helbig I, Afawi Z, Pendziwiat M, Abu-Rachma J, et al. Rare Noncoding Mutations Extend the Mutational Spectrum in the PGAP3 Subtype of Hyperphosphatasia with Mental Retardation Syndrome. Hum Mutat. 2016;37(8):737–44.

8. Tarutani M, Itami S, Okabe M, Ikawa M, Tezuka T, Yoshikawa K, et al. Tissuespecific knockout of the mouse Pig-a gene reveals important roles for GPI-anchored proteins in skin development. Proc Natl Acad Sci U S A. 1997;94(14):7400–5.

9. Kawagoe K, Kitamura D, Okabe M, Taniuchi I, Ikawa M, Watanabe T, et al. Glycosylphosphatidylinositol-anchor-deficient mice: implications for clonal dominance of mutant cells in paroxysmal nocturnal hemoglobinuria. Blood. May 1996;87(9):3600–6.

10. Lukacs M, Blizzard LE, Stottmann RW. CNS glycosylphosphatidylinositol deficiency results in delayed white matter development, ataxia and premature death in a novel mouse model. Hum Mol Genet. 2020;29(7):1205–17.

11. Ahrens MJ, Li Y, Jiang H, Dudley AT. Convergent extension movements in growth plate chondrocytes require gpi-anchored cell surface proteins. Development. 2009;136(20):3463–74.

12. Visconte V, Raghavachari N, Liu D, Keyvanfar K, Desierto MJ, Chen J, et al. Phenotypic and functional characterization of a mouse model of targeted Pig-a deletion in hematopoietic cells. Haematologica. 2010;95(2):214–23.

13. Lukacs M, Roberts T, Chatuverdi P, Stottmann RW. Glycosylphosphatidylinositol biosynthesis and remodeling are required for neural tube closure, heart development, and cranial neural crest cell survival. Elife. 2019;8:1–30.

14. McKean DM, Niswander L. Defects in GPI biosynthesis perturb Cripto signaling during forebrain development in two new mouse models of holoprosencephaly. Biol Open. 2012;1(9):874–83.

15. Krawitz PM, Schweiger MR, Rödelsperger C, Marcelis C, Kölsch U, Meisel C, et al. Identity-by-descent filtering of exome sequence data identifies PIGV mutations in hyperphosphatasia mental retardation syndrome. Nat Genet [Internet]. October 2010;42(10):827–9. Available at: http://dx.doi.org/10.1038/ng.653

16. Ji YK, Hong Y, Ashida H, Shishioh N, Murakami Y, Morita YS, et al. PIG-V involved in transferring the second mannose in glycosylphosphatidylinositol. J Biol Chem. 2005;280(10):9489–97.

17. Horn D, Krawitz P, Mannhardt A, Korenke GC, Meinecke P. Hyperphosphatasia-mental retardation syndrome due to PIGV mutations: Expanded clinical spectrum. Am J Med Genet Part A. 2011;155(8):1917–22.

18. Horn D, Wieczorek D, Metcalfe K, Baric I, Paležac L, Cuk M, et al. Delineation of PIGV mutation spectrum and associated phenotypes in hyperphosphatasia with mental retardation syndrome. Eur J Hum Genet [Internet]. 2014;22(6):762–7. Available at: http://eutils.ncbi.nlm.nih.gov/entrez/eutils/elink.fcgi?dbfrom=pubmed&id=24129430&retmode=ref&cmd=prlinks papers2://publication/doi/10.1038/ejhg.2013.241

19. Knaus A, Pantel JT, Pendziwiat M, Hajjir N, Zhao M, Hsieh TC, et al. Characterization of glycosylphosphatidylinositol biosynthesis defects by clinical features, flow cytometry, and automated image analysis. Genome Med. 2018;10(1):1–13.

20. Sakaguchi T, Žigman T, Ramadža DP, Omerza L, Pušeljić S, Hrvaćanin ZE, et al. A novel PGAP3 mutation in a Croatian boy with brachytelephalangy and a thin corpus callosum. Hum Genome Var. 2018;5(November 2017):2–5.

21. Jirkof P. Burrowing and nest building behavior as indicators of well-being in mice. J Neurosci Methods [Internet]. 2014;234:139–46. Available at: http://dx.doi.org/10.1016/j.jneumeth.2014.02.001

22. Yuan X, Li Z, Baines AC, Gavriilaki E, Ye Z, Wen Z, et al. A hypomorphic PIGA gene mutation causes severe defects in neuron development and susceptibility to complement-mediated toxicity in a human iPSC model. PLoS One. 2017;12(4):1–19.

23. Van Loo KMJ, Rummel CK, Pitsch J, Müller JA, Bikbaev AF, Martinez-Chavez E, et al. Calcium channel subunit α2σ4 is regulated by early growth response 1 and facilitates epileptogenesis. J Neurosci. 2019;39(17):3175–87.

24. Daniele Mattei, Andranik Ivanov, Marc van Oostrum, Stanislav Pantelyushin, Juliet Richetto, Flavia Mueller, Michal Matheus Beffinger, Linda Schellhammer, Johannes vom Berg, Bernd Wollscheid, Dieter Beule, Rosa Chiara Paolicelli UM. Enzymatic dissociation induces transcriptional and proteotype bias in brain cell populations. Vol 53, Dk. 2020. 1689–1699 p.

25. Wang JY, Yu IS, Huang CC, Chen CY, Wang WP, Lin SW, et al. Sun1 deficiency leads to cerebellar ataxia in mice. DMM Dis Model Mech. 2015;8(8):957–67.

26. Kojic M, Gaik M, Kiska B, Salerno-Kochan A, Hunt S, Tedoldi A, et al. Elongator mutation in mice induces neurodegeneration and ataxia-like behavior. Nat Commun [Internet]. 2018;9(1). Available at: http://dx.doi.org/10.1038/s41467-018-05765-6

27. Kalueff A V., Stewart AM, Song C, Berridge KC, Graybiel AM, Fentress JC. Neurobiology of rodent self-grooming and its value for translational neuroscience. Nat Rev Neurosci [Internet]. 2016;17(1):45–59. Available at: http://dx.doi.org/10.1038/nrn.2015.8

28. Silverman JL, Yang M, Lord C, Crawley JN. Behavioural phenotyping assays for mouse models of autism. Nat Rev Neurosci [Internet]. 2010;11(7):490–502. Available at: http://dx.doi.org/10.1038/nrn2851

29. Ledergerber D, Moser EI. Memory Retrieval: Taking the Route via Subiculum. Curr Biol. 2017;27(22):R1225–7.

30. Hervé JC, Bourmeyster N. Rho GTPases at the crossroad of signaling networks in mammals. Small GTPases. 2015;6(2):43–8.

31. Parri M, Buricchi F, Giannoni E, Grimaldi G, Mello T, Raugei G, et al. EphrinA1 activates a Src/focal adhesion kinase-mediated motility response leading to rho-dependent actino/myosin contractility. J Biol Chem. 2007;282(27):19619–28.

32. Eisenhaber B, Bork P, Eisenhaber F. Post-translational GPI lipid anchor modification of proteins in kingdoms of life: Analysis of protein sequence data from complete genomes. Protein Eng. 2001;14(1):17–25.

33. Kelly RJ, Hill A, Arnold LM, Brooksbank GL, Richards SJ, Mitchell LD, et al. Longterm treatment with eculizumab in paroxysmal nocturnal hemoglobinuria: sustained efficacy and improved survival CME article Long-term treatment with eculizumab in paroxysmal nocturnal hemoglobinuria: sustained efficacy and improved survival. Blood. 2011;117(25):6786–92.

34. Szklarczyk D, Franceschini A, Wyder S, Forslund K, Heller D, Huerta-Cepas J, et al. STRING v10: Protein-protein interaction networks, integrated over the tree of life. Nucleic Acids Res. 2015;43(D1):D447–52.

35. Harbott LK, Nobes CD. A key role for Abl family kinases in EphA receptor-mediated growth cone collapse. Mol Cell Neurosci. 2005;30(1):1–11.

36. Tuomisto L, Lozeva V, Valjakka A, Lecklin A. Modifying effects of histamine on circadian rhythms and neuronal excitability. Behav Brain Res. 2001;124(2):129–35.

37. Thakkar MM. Histamine in the regulation of wakefulness. Sleep Med Rev [Internet]. 2011;15(1):65–74. Available at: http://dx.doi.org/10.1016/j.smrv.2010.06.004

38. Urade Y, Hayaishi O. Prostaglandin D2 and sleep/wake regulation. Sleep Med Rev [Internet]. 2011;15(6):411–8. Available at: http://dx.doi.org/10.1016/j.smrv.2011.08.003

39. Ram A, Pandey HP, Matsumura H, Kasahara-Orita K, Nakajima T, Takahata R, et al. CSF levels of prostaglandins, especially the level of prostaglandin D2, are correlated with increasing propensity towards sleep in rats. Brain Res. 1997;751(1):81–9.

40. Eakin GS, Hadjantonakis A. Production of chimeras by aggregation of embryonic stem cells with diploid or tetraploid mouse embryos. 2006;1(3).

41. R Core Team (2019). R: A language and environment for statistical computing. R Foundation for Statistical Computing, Vienna, Austria. URL https://www.R-project.org/.]

