## Supplementary Information (SI)-Appendix for "A CRISPR-Cas9–engineered mouse model for GPI-anchor deficiency mirrors human phenotypes and exhibits hippocampal synaptic dysfunctions"

### Supplementary Figure 1

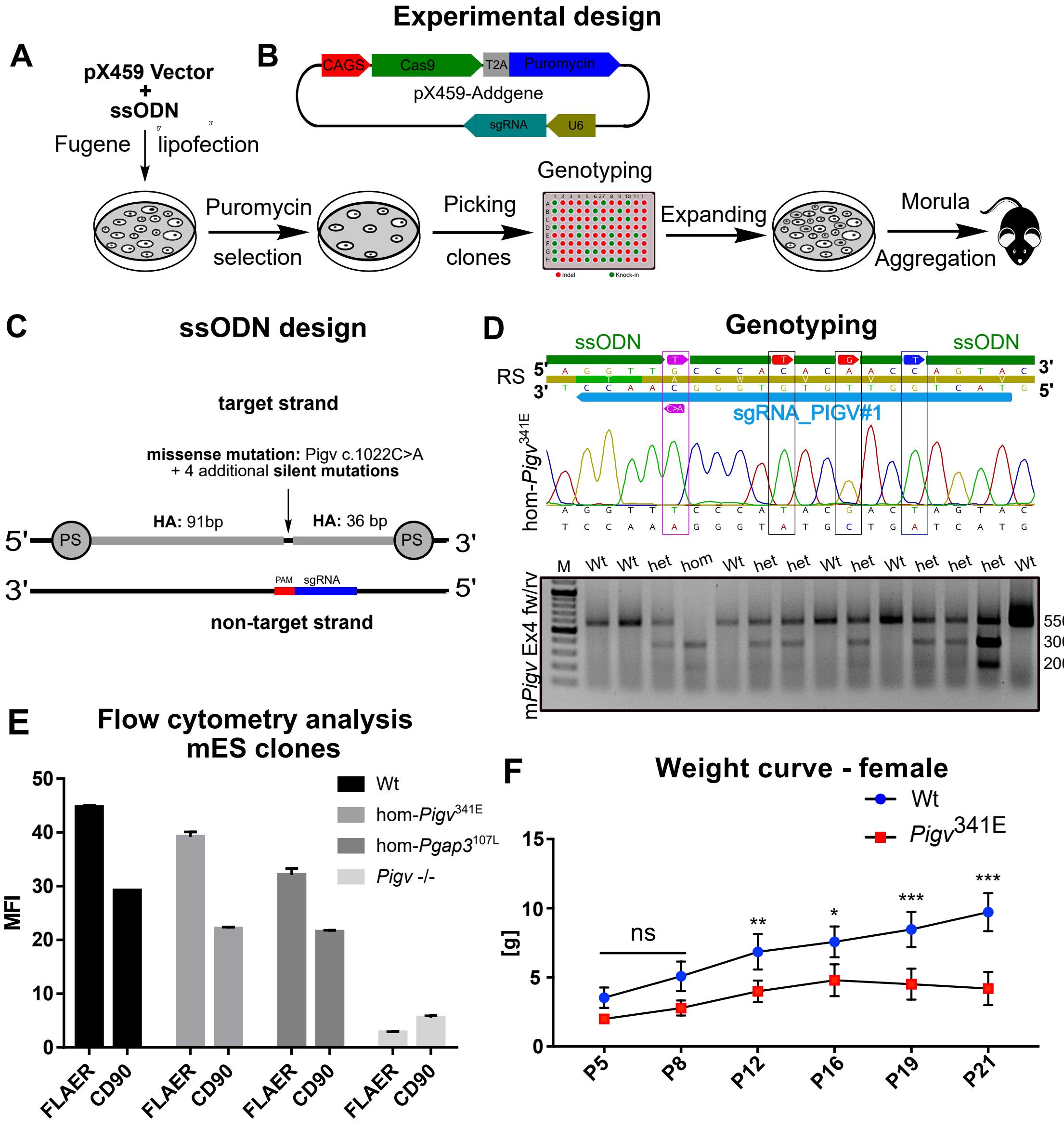


**Generation of the mouse model, analysis of mES clones and weight curve.** (A) Schematic overview of the experimental design from generating the mouse embryonic stem (mES) cell clone to establishment of the mouse model. (B) Schematic overview of the pX459 vector from Addgene. (C) single-stranded oligode-oxynucleotides (ssODNs) were designed with asymmetrical homology arms (HA), highlighted in grey and with phosphorothioate (PS) bonds according to Renaud et al. (2016), Richardson et al. (2016). ssODNs were identical to the target strand. Single-guide RNA (sgRNA) is highlighted in blue. Protospacer adjacent motif (PAM) is marked in red (D) ssODNs contained the nucleotide T (marked in pink) causing the missense mutation c.1022 C>A on the complementary strand and three additional silent mutations (marked in red and blue) that avoid a second cut by Cas9 after homology-directed repair (HDR) has happened. The silent mutation marked in blue generated a restriction site for *BcuI* that was used for genotyping purposes. Sanger sequencing of homozygous *Pigv*$^{341E}$ knock-in clone revealed the stable integration of missense mutation (on the (-) strand C>A, on the (+) strand G>T, causing PIGV p.Ala341Glu) and silent mutations (on the (+) strand C>T, A>G, C>T) at CRISPR-Cas9-induced DSB by HDR. Gel electrophoresis identified genotypes in mice by *BcuI* digest of PCR-amplified products using the primers m*Pigv*_Ex4_fw and m*Pigv*_Ex4_rv. RS: Reference sequence of *Pigv* (E) Flow cytometry analysis of CRISPR-engineered mouse embryonic stem cell (mES) clones: homozygous *Pigv*$^{341E}$ (hom-*Pigv*$^{341E}$), homozygous *Pgap3*$^{107L}$ (hom-*Pgap3*$^{107L}$) +/+, homozygous *Pigv* knock-out (*Pigv* -/-) revealed decreased expression in FLAER and GPI-linked CD90 as compared to wildtype (wt), MFI: Mean fluorescence intensity. (F) Female *Pigv*$^{341E}$ mice reveal a reduced weight from postnatal day (P) 1 to 21. Animals used for the weight curve: Wt(female n=4) *Pigv*$^{341E}$(female n= 3). The data from the weight curve was analyzed by two-way of variance (ANOVA) followed by a Bonferroni's multiple comparisons test. *P<0.05, **P<0.01, ***P<0.001.

### Supplementary Figure 2

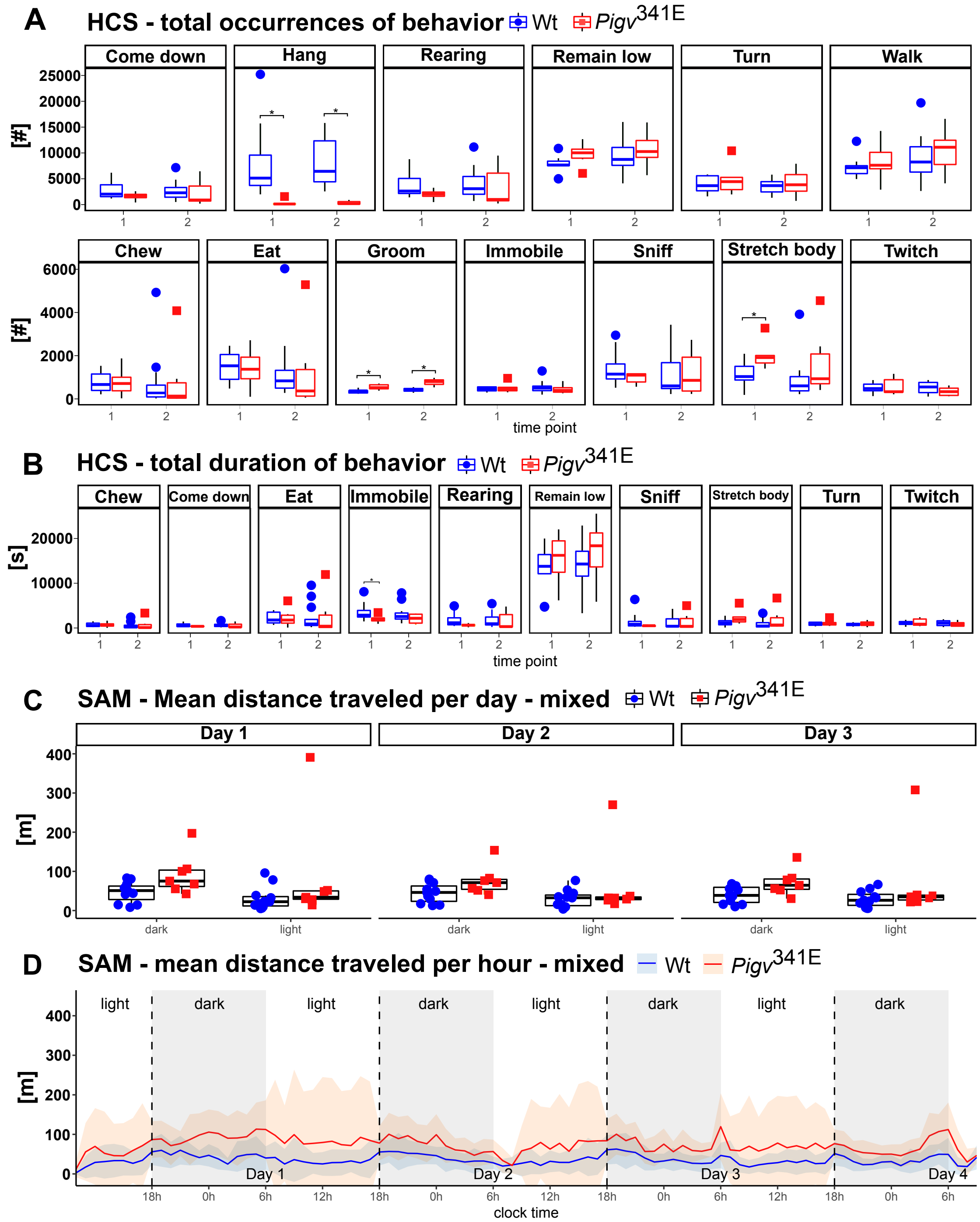


**HCS and SAM (second approach)** (A) Total occurrences of behaviors from the home cage scan (HCS). *Pigv*$^{341E}$ mice revealed a decreased number of hanging and an increased number of grooming at both time points. *Pigv*$^{341E}$ mice showed an increased number of "stretch body" at time point 1. (B) Total duration of behavior of behaviors from the HCS. *Pigv*$^{341E}$ mice showed a decreased "immobile" behavior at time point 1. (C, D) In the social activity monitor (SAM), no significant differences in mean distance traveled per day and per hour were observed between genotypes when held in mixed genotypes. *Pigv*$^{341E}$=homozygous for Pigv p.Ala341Glu, wt=wild-type. Animals used for the HCS were at time point 1 (tp1) 8 weeks old, at time point 2 (tp2) 16 weeks old: Wt(female n=8, male n=4) *Pigv*$^{341E}$(female n= 4, male n=2). Animals used for SAM: Wt(female n=4, male n=6) *Pigv*$^{341E}$(female n= 2, male n=5). The data from the HCS (total occurrences, total duration) was analyzed with Wilcoxon rank sum test (non parametric). The data from the SAM (distance traveled per hour) was analyzed with generalized linear mixed-effects models (glmm) using a Markov chain Monte Carlo (MCMC).*P<0.05.

### Supplementary Figure 3

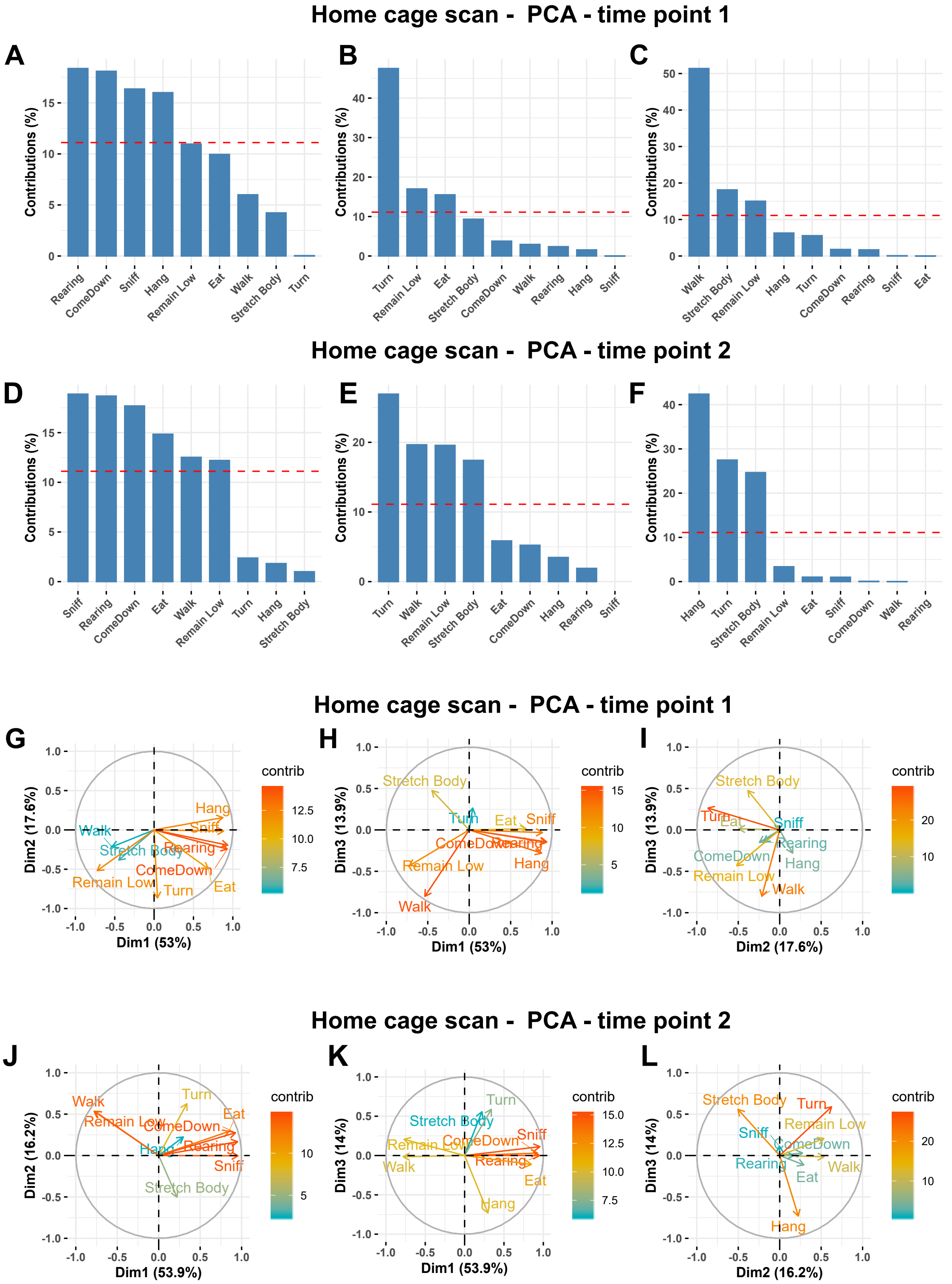


**HCS (Home cage scan) – principal component analysis (PCA).** (A-F) contribution of original variables (occurrences of behavior) to the first three components of the PCA for each time point. (G-L) 2D Representation of the coefficient of all variables along PC1 and PC2 (left), PC1 and PC3 (middle) and PC2 and PC3 (right) for each time point.

### Supplementary Figure 4

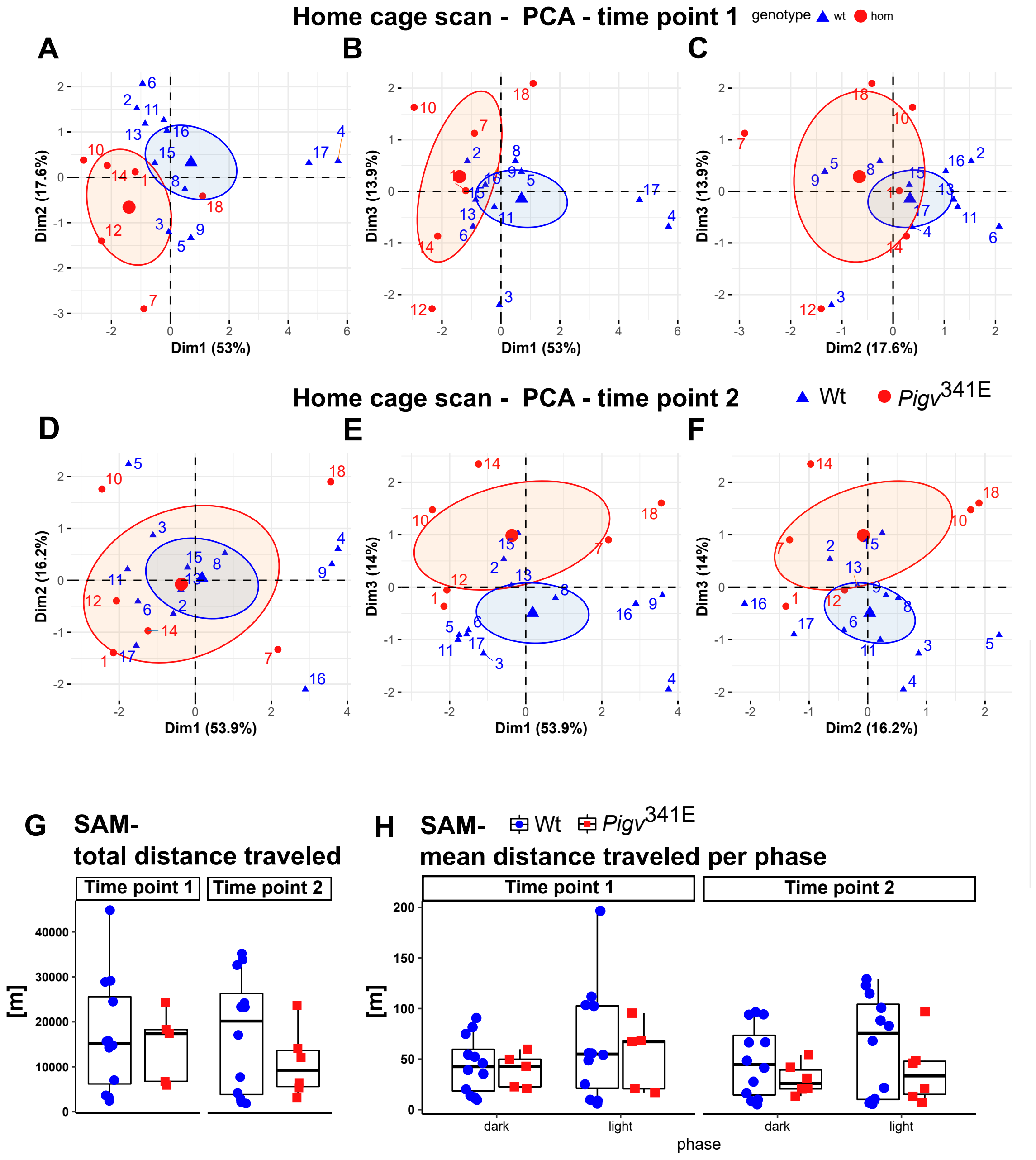


**HCS (Home cage scan) – principal component analysis (PCA), SAM (first approach).** (A-F) Group separation and individual contribution to PC1 and PC2 (left), PC1 and PC3 (middle) and PC2 and PC3 (right). Mean (bigger symbol) and confidence interval (ellipse) of each group are shown. Genotypes differ significantly in dimension 1 at time point 1 and in dimension 3 in time point 2 (G-H) Social activity monitor (SAM) showed no significant differences in total distance traveled and distance traveled per phase between genotypes when held in mixed genotypes.

*Pigv*$^{341E}$=homozygous for Pigv p.Ala341Glu, wt=wild-type. Animals used for the HCS were at time point 1 (tp1) 8 weeks old, at time point 2 (tp2) 16 weeks old: Wt(female n=8, male n=4) *Pigv*$^{341E}$(female n= 4, male n=2).Animals used for the SAM were at time point 1 (tp1) 9 weeks old: Wt(female n=8, male n=4), *Pigv*$^{341E}$(female n= 3, male n=2). Animals were at time point 2 (tp2) 17 weeks old: Wt(female n=8, male n=4), *Pigv*$^{341E}$(female n= 4, male n=2). Data from PCA and SAM (total distance traveled, mean distance traveled per phase) was analyzed with a Wilcoxon rank sum test. *P<0.05, **P<0.01, ***<P0.001.

### Supplementary Figure 5

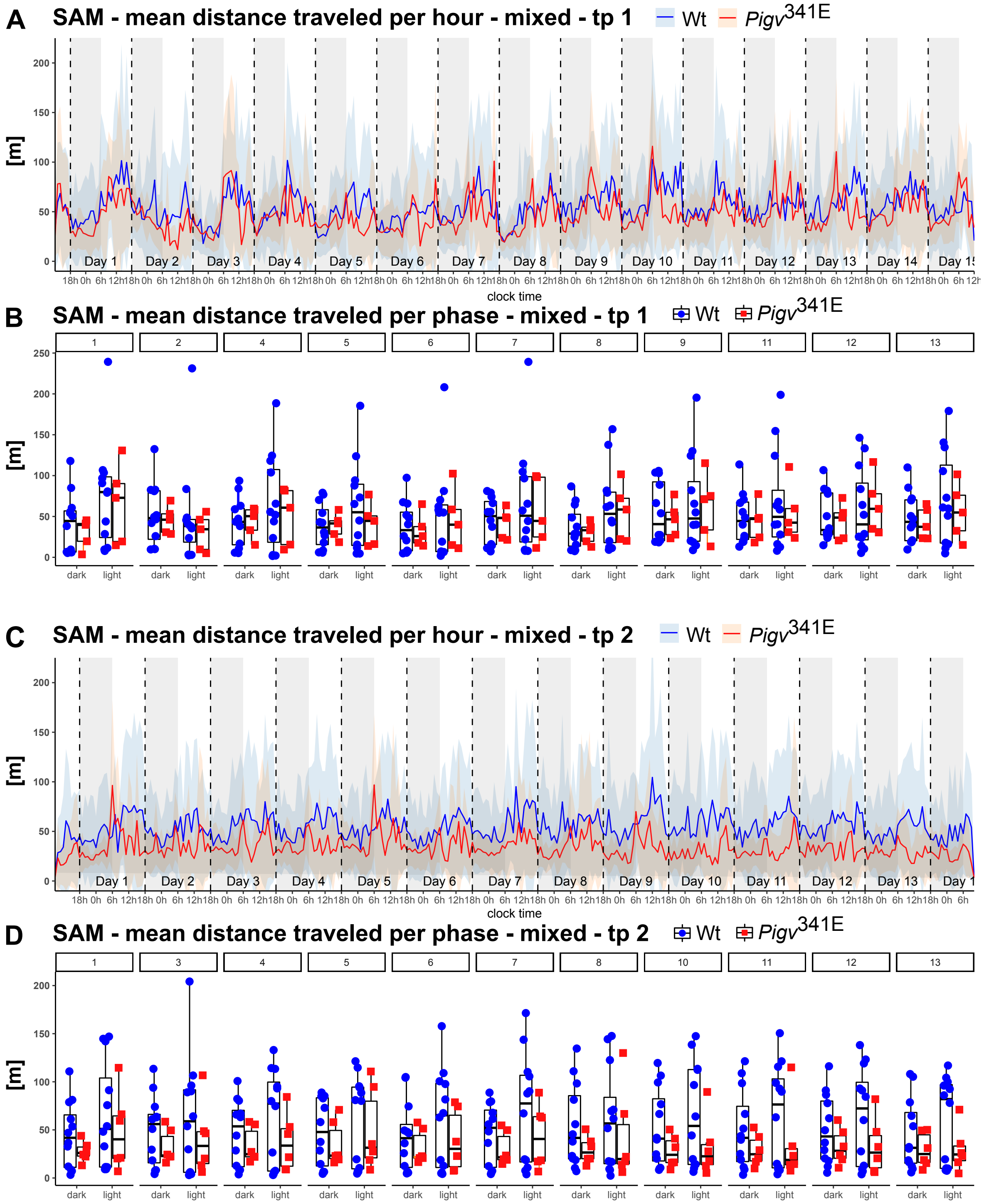


**SAM (first approach).** (A, C) In the social activity monitor (SAM), no significant differences were observed in distance traveled per hour at both time points between genotypes when animals were held in mixed genotypes. Both genotypes were significantly more active during the light cycle than during the dark cycle at both time points. (B, D) No significant differences were observed in distance traveled per phase at both time points between genotypes when animals were held in mixed genotypes. *Pigv*$^{341E}$=homozygous for Pigv p.Ala341Glu, wt=wild-type. Animals used for the SAM were at time point 1 (tp1) 9 weeks old: Wt(female n=8, male n=4), *Pigv*$^{341E}$(female n= 3, male n=2). At time point 2 (tp2) animals were 17 weeks old: Wt(female n=8, male n=4), *Pigv*$^{341E}$(female n= 4, male n=2). Data from the SAM (mean distance per hour) was analyzed with a generalized linear mixed-effects models (glmm) using a Markov chain Monte Carlo (MCMC). *P<0.05, **P<0.01, ***P<0.001.

**Supplementary Figure 6**

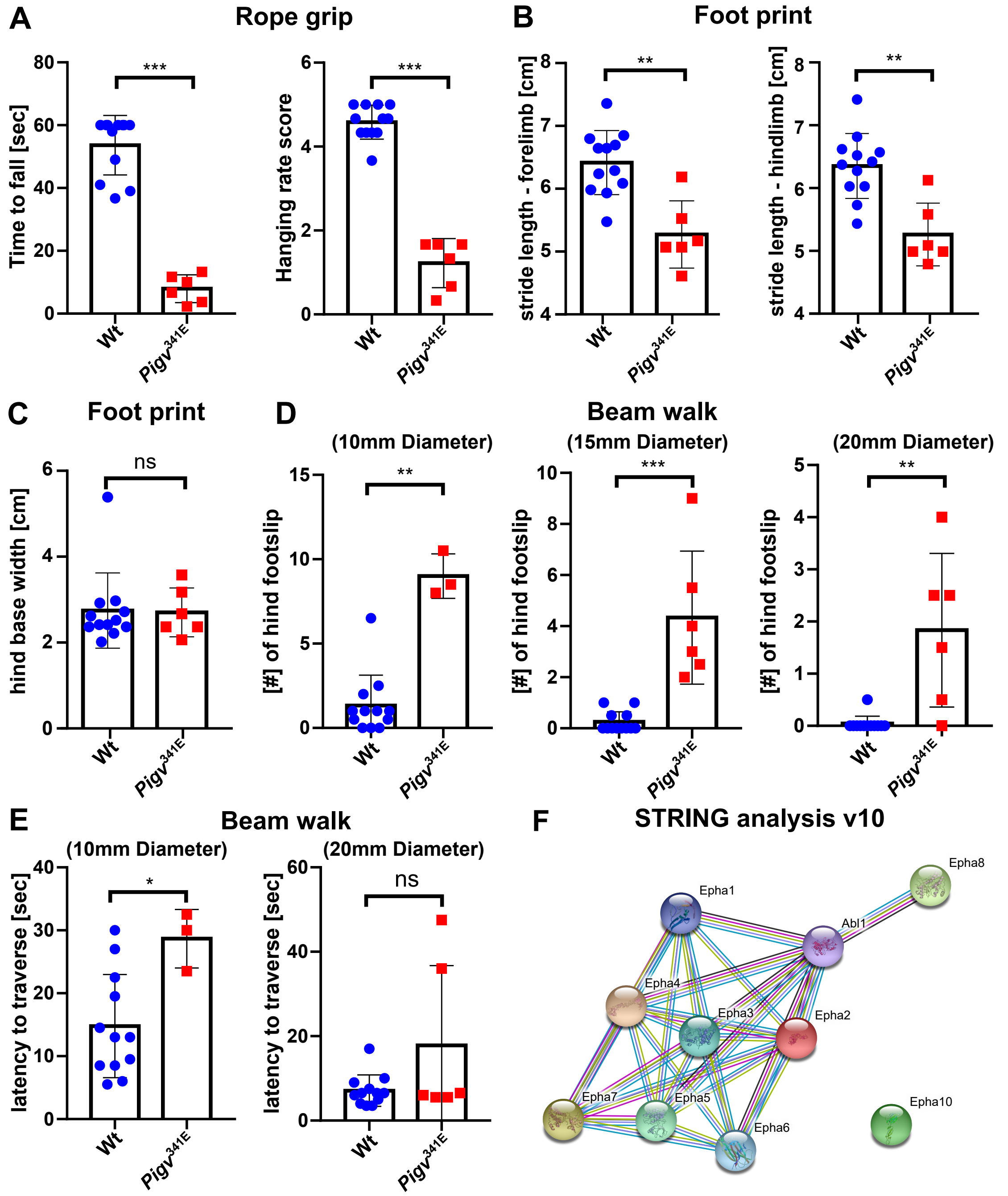


***Pigv*$^{341E}$ mice show a motoric phenotype and STRING analysis** (A) In the rope grip test, *Pigv*$^{341E}$ mice showed a decreased climbing performance displayed by a decreased time to fall off the rope (left graph) and a decreased hanging score (right graph). (B,C) In the foot print test, *Pigv*$^{341E}$ mice exhibited no significant differences to wild-type in hind base width but showed a decreased forelimb and hindlimb stride length (FL-SL, HL-SL). (D) *Pigv*$^{341E}$ mice had deficits in motor coordination displayed by an increased number of hind footslip when walking on beams with different diameters (10-20 mm diameter). (E) *Pigv*$^{341E}$ mice showed an increased latency to traverse a beam with a diameter of 10 mm (left graph) but revealed no significant difference to wild-type mice traversing a beam of 20 mm (right graph). (F) STRING analysis v10 confirmed protein-protein interaction between EphA-receptors and Abl1. Turquoise string: from curated databases, pink string: experimentally determined, yellow string: textmining, black string: co-expression, lila string: protein homology.

*Pigv*$^{341E}$=homozygous for Pigv p.Ala341Glu, wt=wild-type. Wt(female n=8, male n=4), *Pigv*$^{341E}$ (female n= 4, male n=2). Animals were 7 weeks old. The data was analyzed with a non-parametric t-test (Mann-Whitney). *P<0.05, **P<0.01, ***P<0.001

**Supplementary Figure 7**

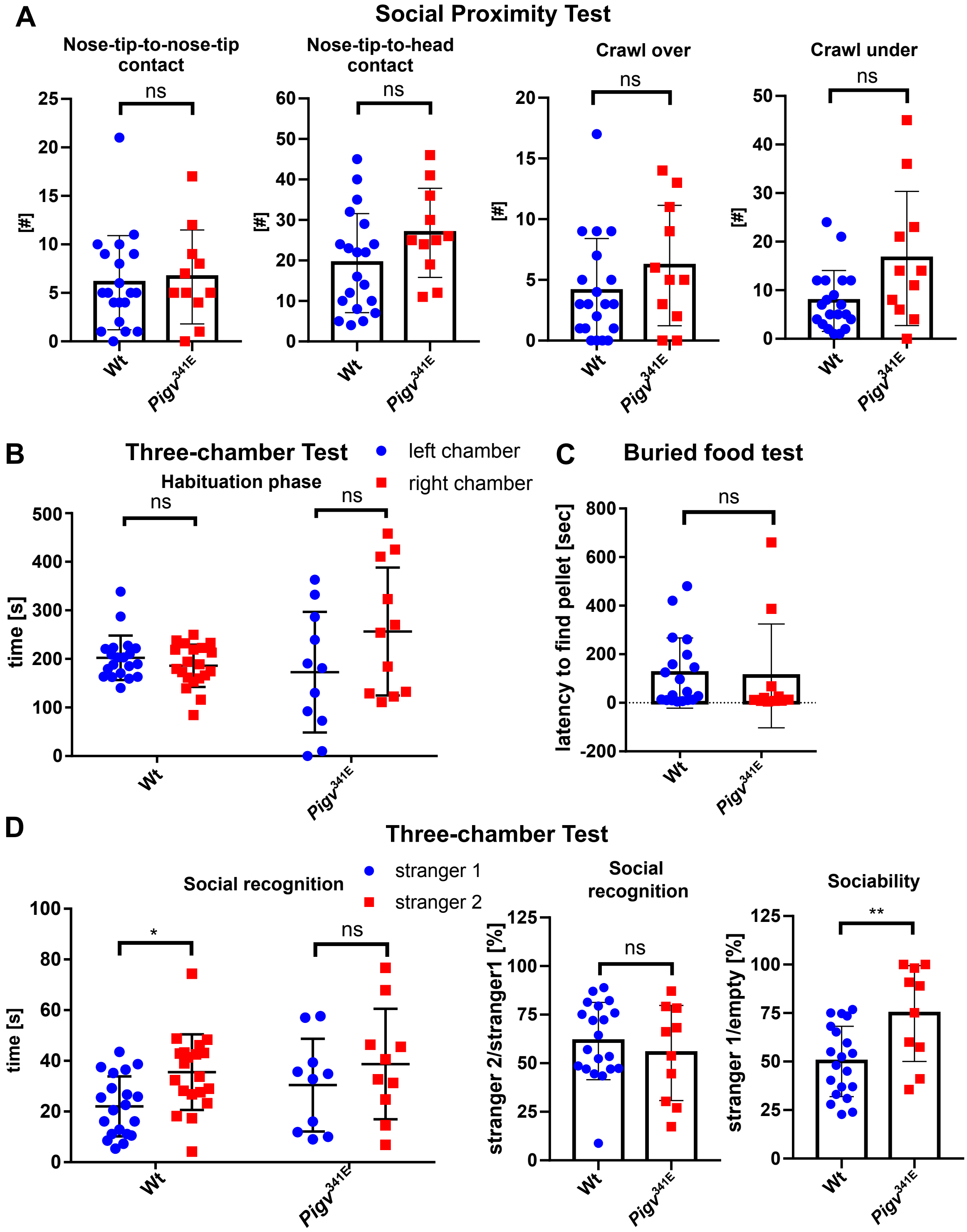


***Pigv*$^{341E}$ mice exhibit a social phenotype and no defects in olfaction** (A) In the social proximity test, *Pigv*$^{341E}$ mice revealed no significant differences to wild-type in numbers of nose-tip-to-nose-tip, nose-tip-to-head contact, "crawl over" and "crawl under" behavior when interacting with the stranger mouse. (B) During the habituation phase *Pigv*$^{341E}$ and wild-type mice did not show any preferences for the left and right chamber. (C) In the buried food test, no differences were observed in olfaction between genotypes. (D) In contrast to wild-type, *Pigv*$^{341E}$ mice showed no significant difference in spending time with stranger 1 and 2 in the three-chamber test (first 5 minutes) (left graph) indicating impairment in social recognition. However, ratio between stranger 2 and 1 (middle graph) did not show any differences between genotypes in the three-chamber test (first 5 minutes). The ratio between stranger 1 and empty cage was increased (right graph) in *Pigv*$^{341E}$ mice indicating an enhanced social approach behavior in the three-chamber test (first 5 minutes). *Pigv*$^{341E}$=homozygous for Pigv p.Ala341Glu, wt=wild-type. Animals used for the three-chamber test: Wt(female n=9, male n=11), *Pigv*$^{341E}$(female n=4, male n=6). Animals used for the social proximity test and buried food test: Wt(female n=9, male n=11), *Pigv*$^{341E}$(female n=4, male n=7). Stranger mice had the same age and sex as testing mice. The data from the social proximity test, three-chamber test (ratio strainger2/1, stranger1/empty) and buried food test was analyzed with a non-parametric t-test (Mann-Whitney). The data from the three-chamber test (habituation phase, social recognition) was analyzed by two-way of variance (ANOVA) followed by a Bonferroni's multiple comparisons test. *P<0.05, **P<0.01, ***P<0.001.

### Supplementary Figure 8

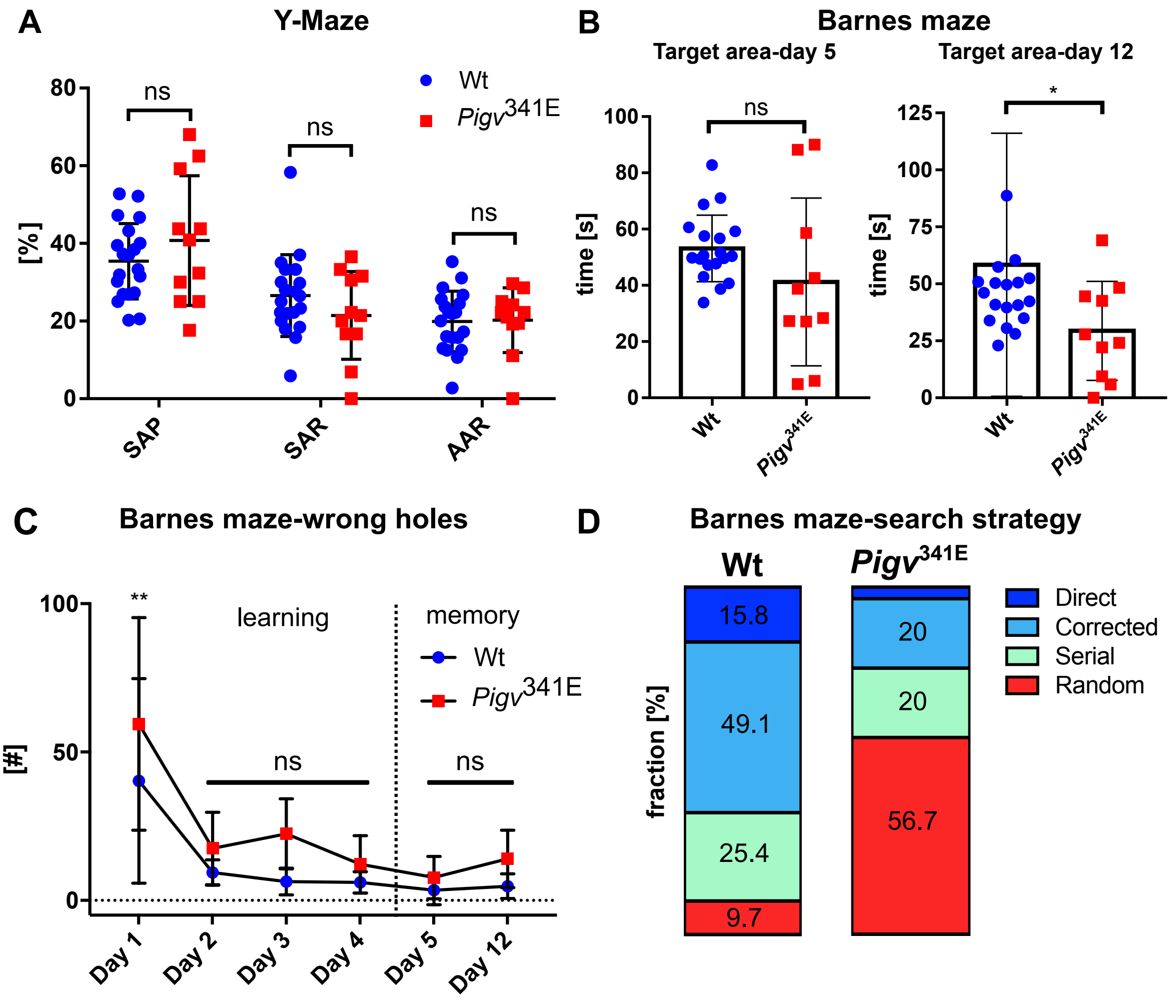


***Pigv*341E mice show cognitive deficits in spatial memory.** (A) In the y-maze, *Pigv*341E mice showed no alterations in spontaneous alternation performance (SAP), same arm returns (SAR) and alternate arm returns (AAR) and thus are not impaired in short-term working memory. (B) *Pigv*341E mice spent less time in the target quadrant of the Barnes maze at day 12 (long-term memory). At day 5 (short-term memory) there was no significant difference between genotypes. (C) In the Barnes maze, *Pigv*341E mice showed no significant differences in the number of wrong holes to wild-type during day 2-12. (D) Fractions of search strategies during the Barnes maze test that *Pigv*341E mice and wild-type used in order to find the correct exit hole were shifted in *Pigv*341E mice. Direct=direct way to the correct exit hole, corrected=mice targeted first 1-3 wrong holes and went afterwards directly to the correct hole, serial=mice checked all holes in series, random=mice targeted random holes without any order.
*Pigv*341E=homozygous for Pigv p.Ala341Glu, wt=wild-type. Animals used for the y-maze: Wt(female n=9, male n=11), *Pigv*341E(female n=4, male n=7). Animals used for the Barnes maze: Wt(female n=8, male n=11) *Pigv*341E(female n=4, male n=6). The data from the y-maze and the barnes maze (target area day 5 and 12) was analyzed with a non-parametric t-test (Mann-Whitney). The data from the Barnes maze (number of wrong holes) was analyzed by two-way of variance (ANOVA) followed by a Bonferroni's multiple comparisons test. *P<0.05 **P<0.01, ***P<0.001.

**Supplementary Figure 9**

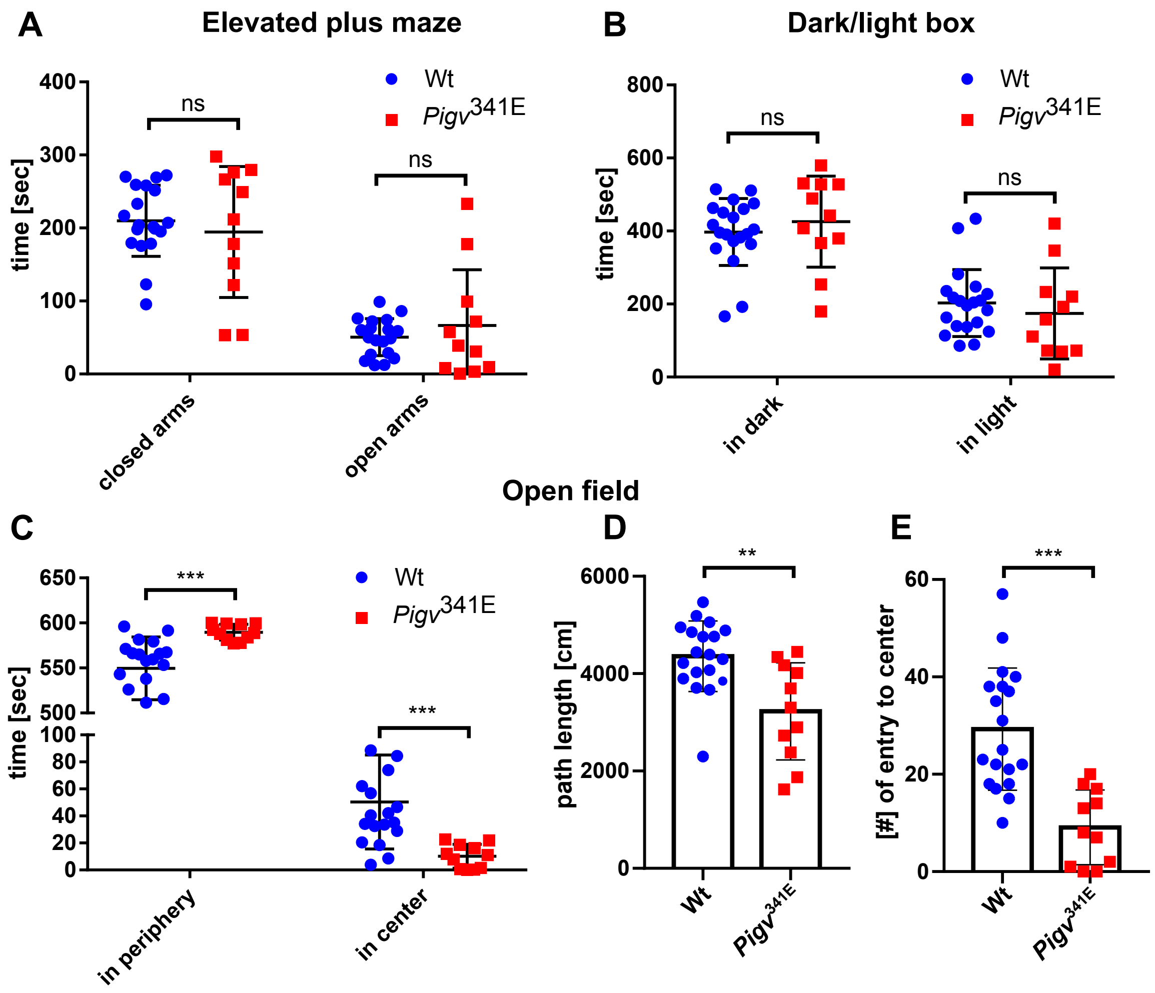


**Affective behavior of *Pigv*[341E] mice.** (A) In the elevated plus maze, *Pigv*[341E] mice showed no significant difference to wild-type in exploration behavior displayed by spending the same time in the open and closed arms. (B) In the dark/light box, *Pigv*[341E] mice exhibited no significant difference to wild-type in exploring the dark and light compartment. (C) In the open field test, *Pigv*[341E] mice spent more time in the periphery than in the center. (D) *Pigv*[341E] mice showed a reduced distance traveled in the open field displayed by a reduced path length. (E) *Pigv*[341E] mice visited less time the center of the open field.

*Pigv*[341E]=homozygous for Pigv p.Ala341Glu, wt=wild-type. Animals used for the the elevated plus maze, dark/light box: Wt(female n=9, male n=11) *Pigv*[341E] (female n=4, male n=7). The data from the open field, elevated plus maze, dark/light box was analyzed with a non-parametric t-test (Mann-Whitney). *P<0.05 **P<0.01, ***P<0.001.

### Supplementary Figure 10

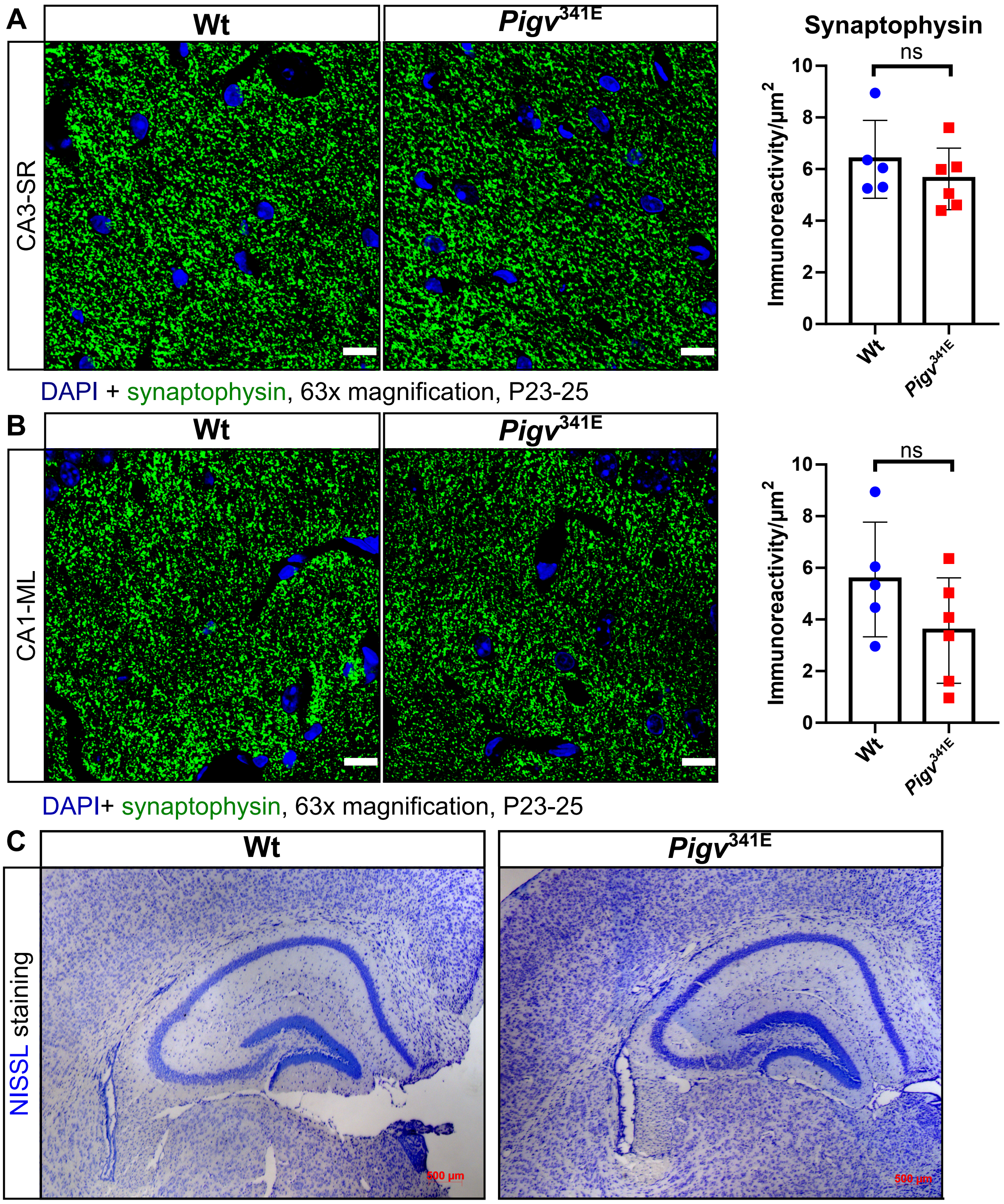


***Pigv*[341E] mice show no morphological abnormalities in the hippocampus** (A-B) Representative image of immunofluorenscence staining showed no difference in synaptophysin immunoreactivity (green) in *Cornu Ammonis* 3 - *Stratum Radiatum* (CA3-SR) and *Cornu Ammonis* 1 – molecular layer of dentate gyrus (CA1-ML) between genotypes (left). 4′,6-Diamidin-2-phenylindol (DAPI) (blue). Images were taken at 63x magnification, scale=10 µm. Synaptophysin immunoreactivity were quantified with Fiji ImageJ and revealed no significantly reduced immunoreactivity/$\mu m^2$ in CA3-SR and CA1-ML between genotypes (right). Animals were 3 weeks old: Wt(female n=3, male n=2) *Pigv*[341E](female n=2, male n=4). (C) Representative images of a Nissl staining revealed no morphological abnormalities in the hippocampus between genotypes. 4x magnification, scale=500 µm. The data from the synaptophysin immunofluorescence staining was analyzed with a parametric student t-test. *P<0.05 **P<0.01, ***P<0.001.

**Supplementary Figure 11**

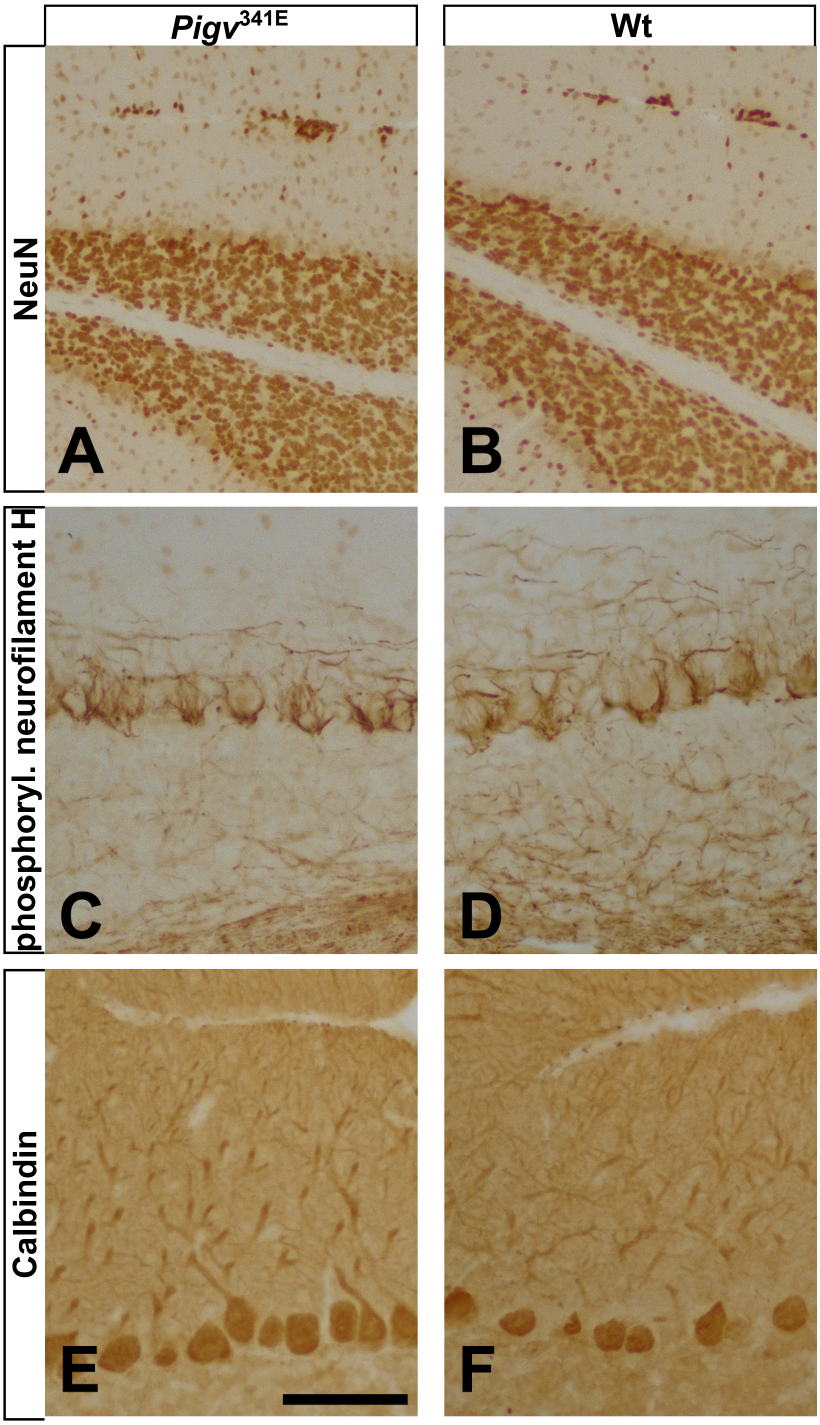


***Pigv*$^{341E}$ mice show no morphological abnormalities in the cerebellum** (A-B) Representative image of immuno-3,3′-Diaminobenzidin (DAB) staining showed no changes of labels granule cells in the granule cell layer (shown are the granule cell layers from lobule 8) between genotypes. A few scattered cells were observed in the molecular layer, and small granule cell ectopias in the meninges separating lobules 8 and 7 (at the top of the panels). Such ectopias are common in C57bl mice, as used here, and no differences in the position or extent of these ectopias could be detected for wild type and Pigv$^{341E}$ mice. (C-D) Phosphorylated neurofilament H outlines axons of basket and stellate cells that converge from the molecular layer (top of the panels) to form the name-giving axonal basket around Purkinje cell perikarya. No differences were observed between genotypes. (E-F) Staining for calbindin D28k reveals regularly arranged. Bar (in panel E) = 80 µm for panels A and B, and 40 µm for panels C-F. Animals were 3 weeks old.

### Supplementary Figure 12

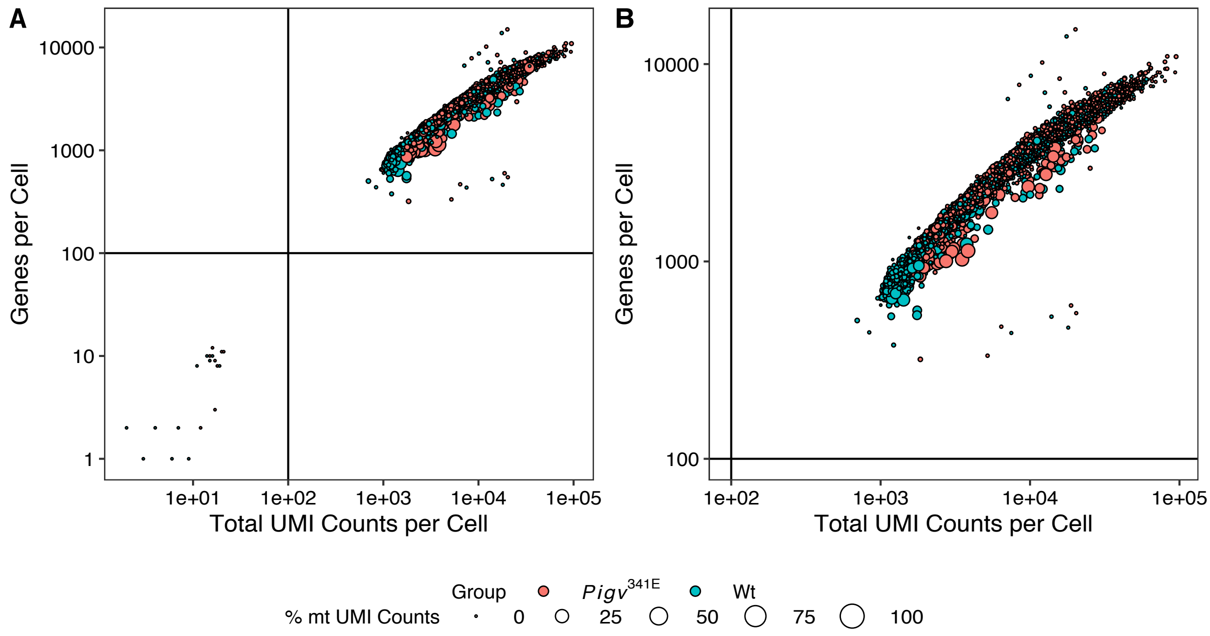


**Filtering removes outlying barcodes.** Keeping only cellular barcodes with more than 100 genes per cell, more than 100 UMI counts per cell and less than 30 % UMI counts mapping to mitochondrial genes from the full preprocessed data set comprising 15,949 barcodes (A) removes 19 outlying barcodes, resulting in the filtered data set comprising 15,930 barcodes (B).

### Supplementary Figure 13

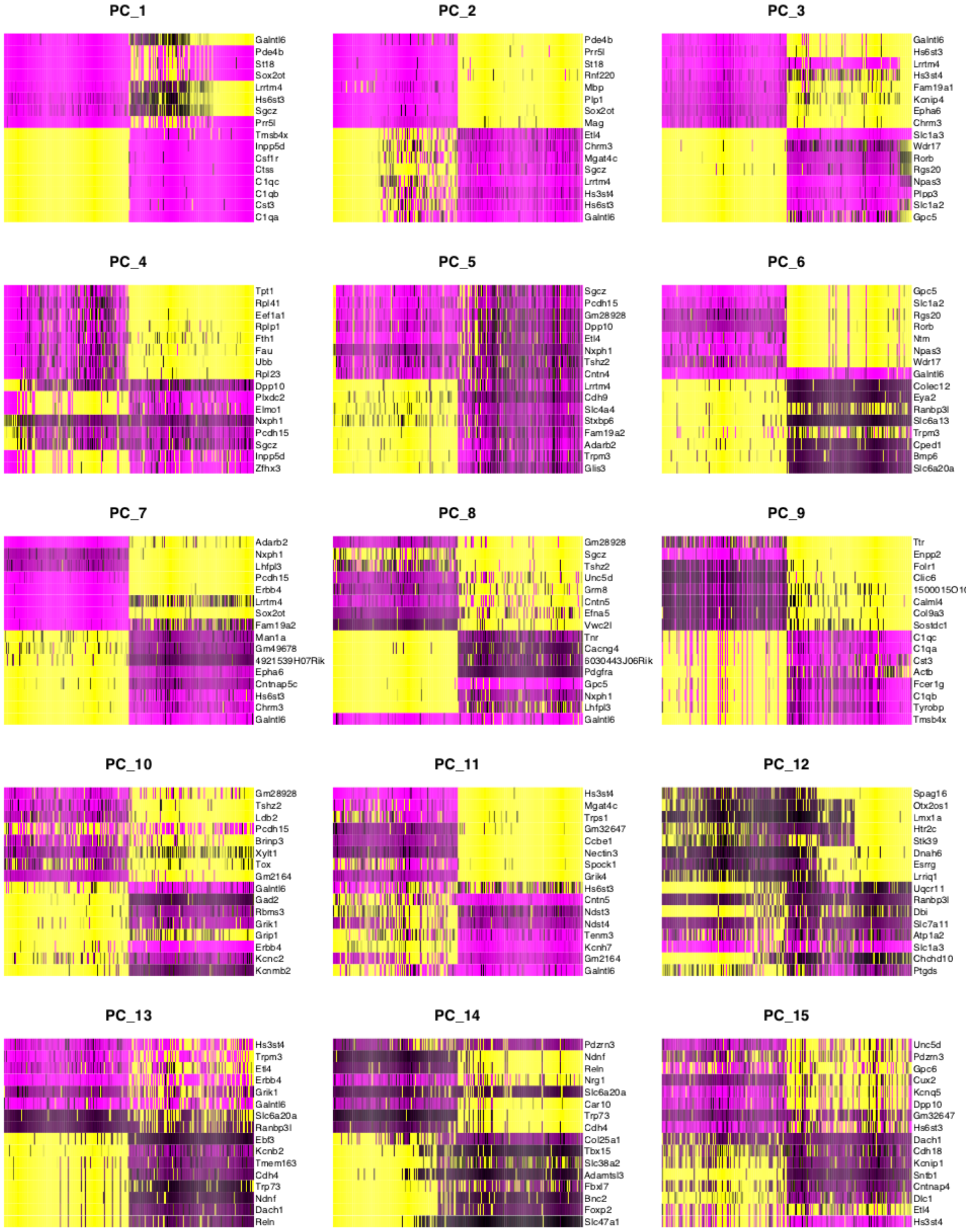


**Dimensional reduction heat maps.** Heat maps on the 15 most influential genes and 250 cells sorted by principal component score for each of the first 15 principal components support the selection of the top 11 principal components for further non-parametric dimensionality reduction and downstream analysis.

### Supplementary Figure 14

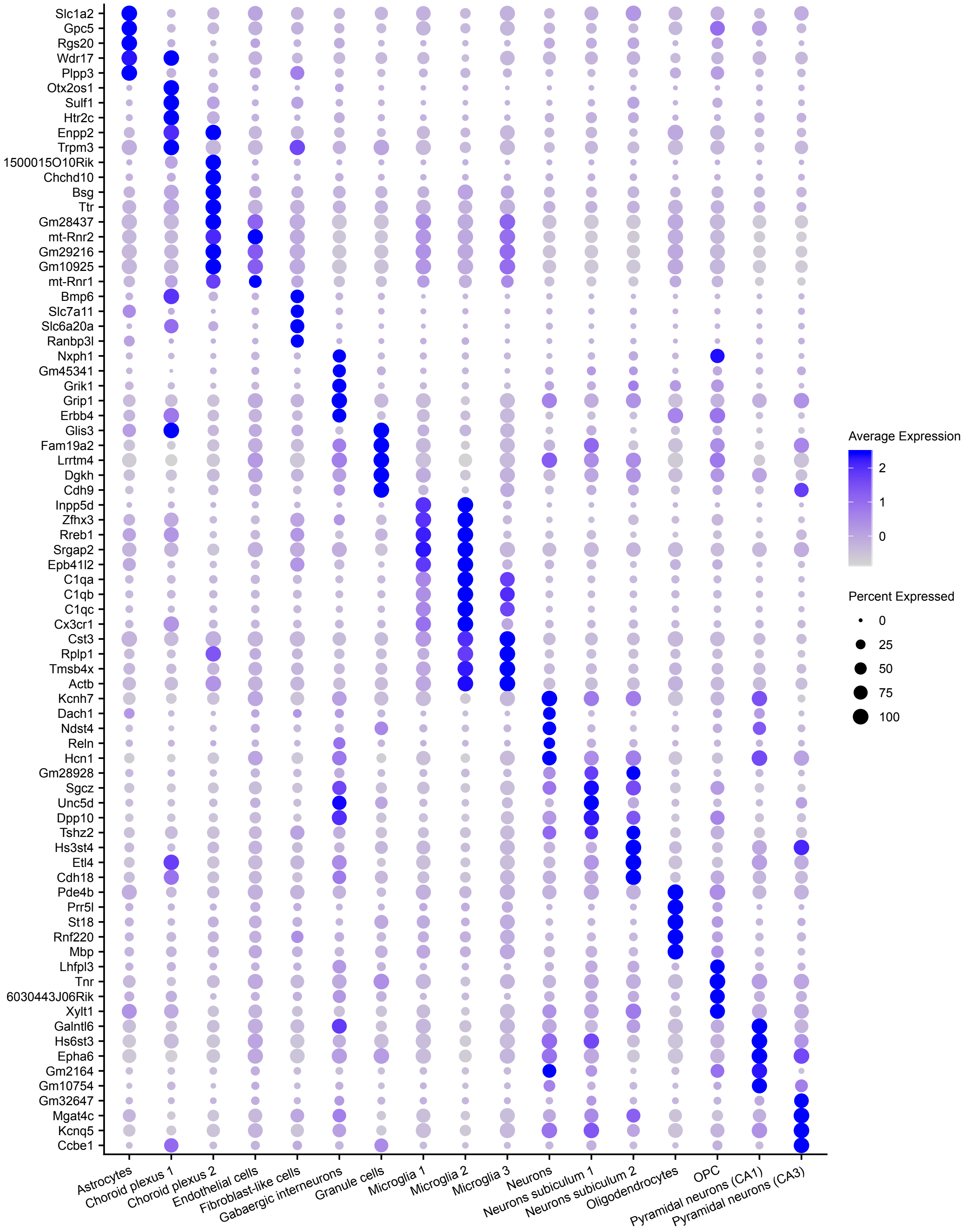


**Expression of marker genes in cellular subgroups.** For each of the 17 cellular subgroups, the 5 differentially expressed genes with highest average fold change in comparison to all remaining cells are depicted. The dot size represents the percentage of cells within a cluster with non-zero expression of the respective gene.

**Supplementary Figure 15**

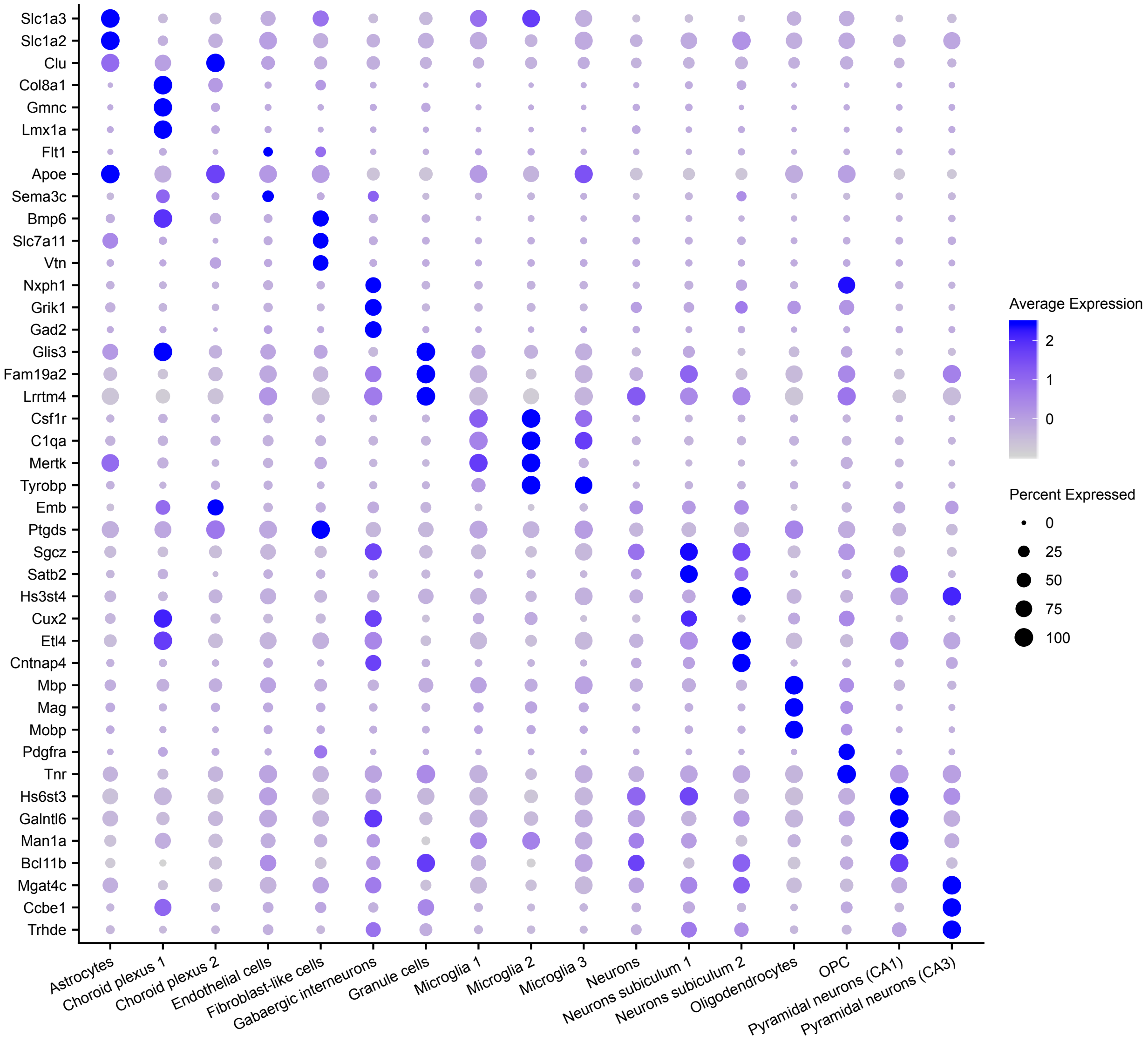


**Expression of selected genes in cellular subgroups.** For each of the 17 cellular subgroups, the average expression of selected genes that supported assignment of cellular subgroup identities are depicted. The dot size represents the percentage of cells within a cluster with non-zero expression of the respective gene.

### Supplementary Figure 16

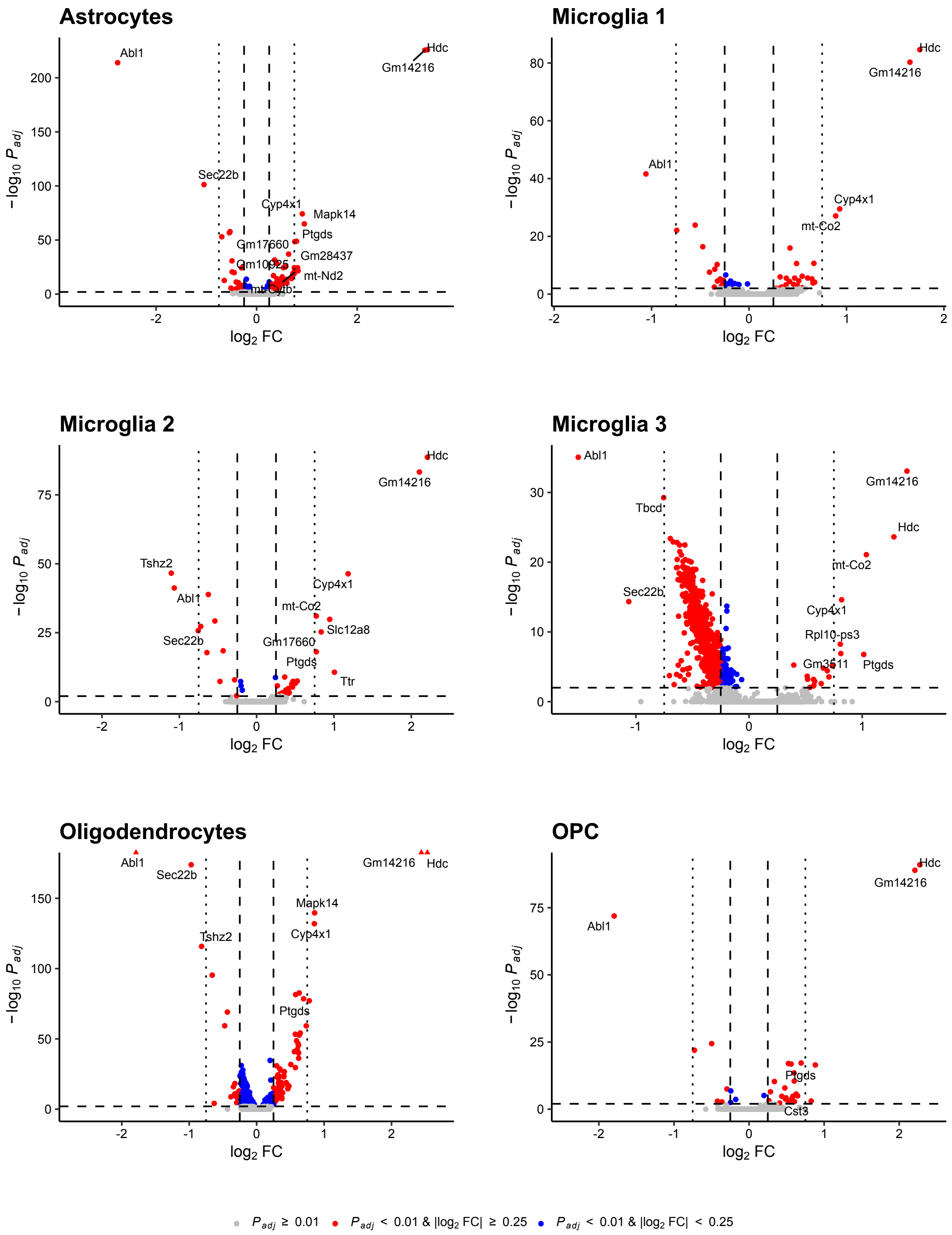


**Volcano plots resulting from comparison of *Pigv*[341E] cells and wild-type cells within glial cellular subgroups.** Dashed horizontal lines are located at an adjusted p-value of 0.01, dashed vertical lines at an absolute $\log_2$ fold change of 0.25 and dotted vertical lines at an absolute $\log_2$ fold change of 0.75. For genes with an adjusted p-value below 0.01 and a minimum absolute $\log_2$ fold change of 0.75, gene symbols are depicted.

### Supplementary Figure 17

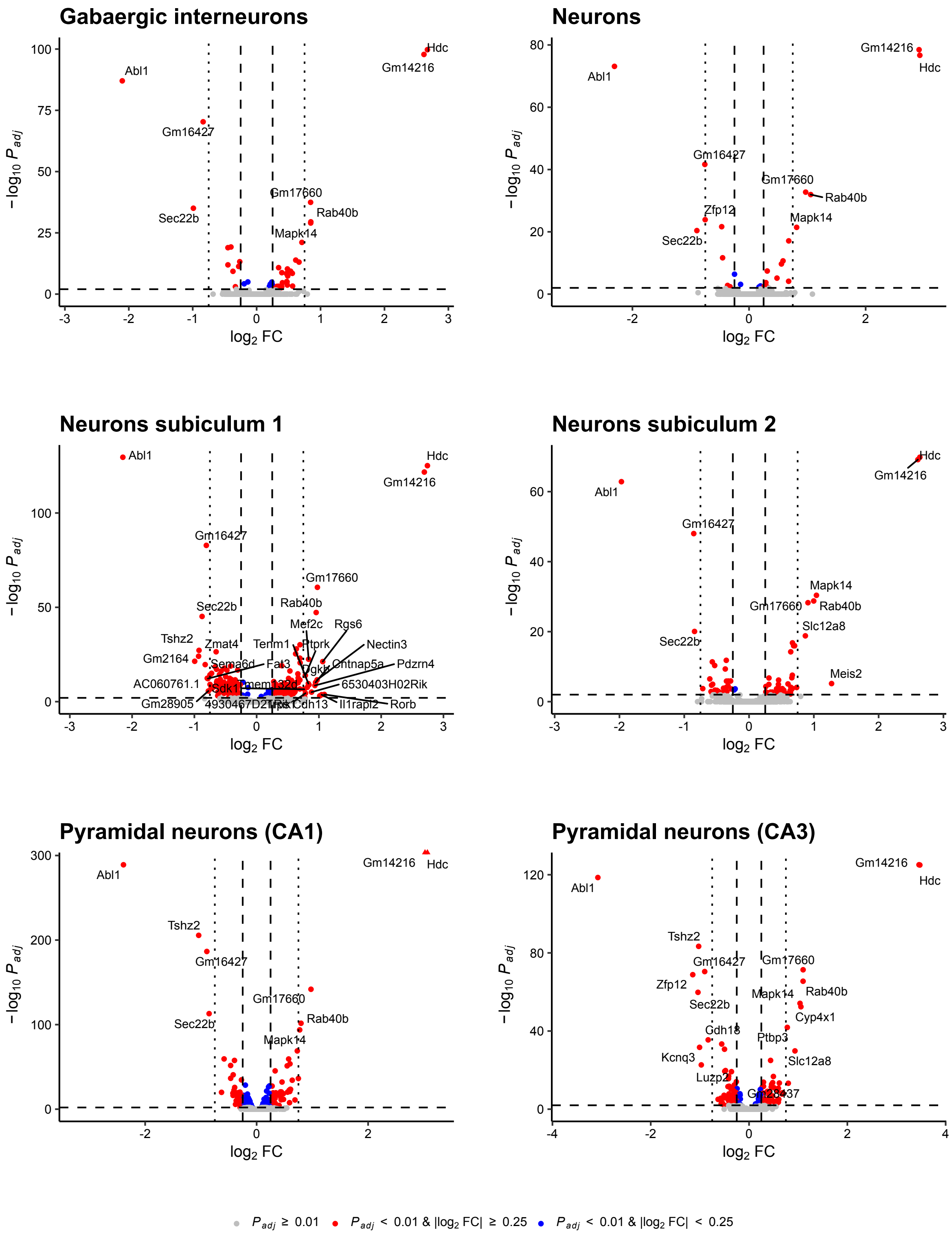


**Volcano plots resulting from comparison of *Pigv*$^{341E}$ cells and wild-type cells within neuronal cellular subgroups.** Dashed horizontal lines are located at an adjusted p-value of 0.01, dashed vertical lines at an absolute $\log_2$ fold change of 0.25 and dotted vertical lines at an absolute $\log_2$ fold change of 0.75. For genes with an adjusted p-value below 0.01 and a minimum absolute $\log_2$ fold change of 0.75, gene symbols are depicted.

### Supplementary Figure 18

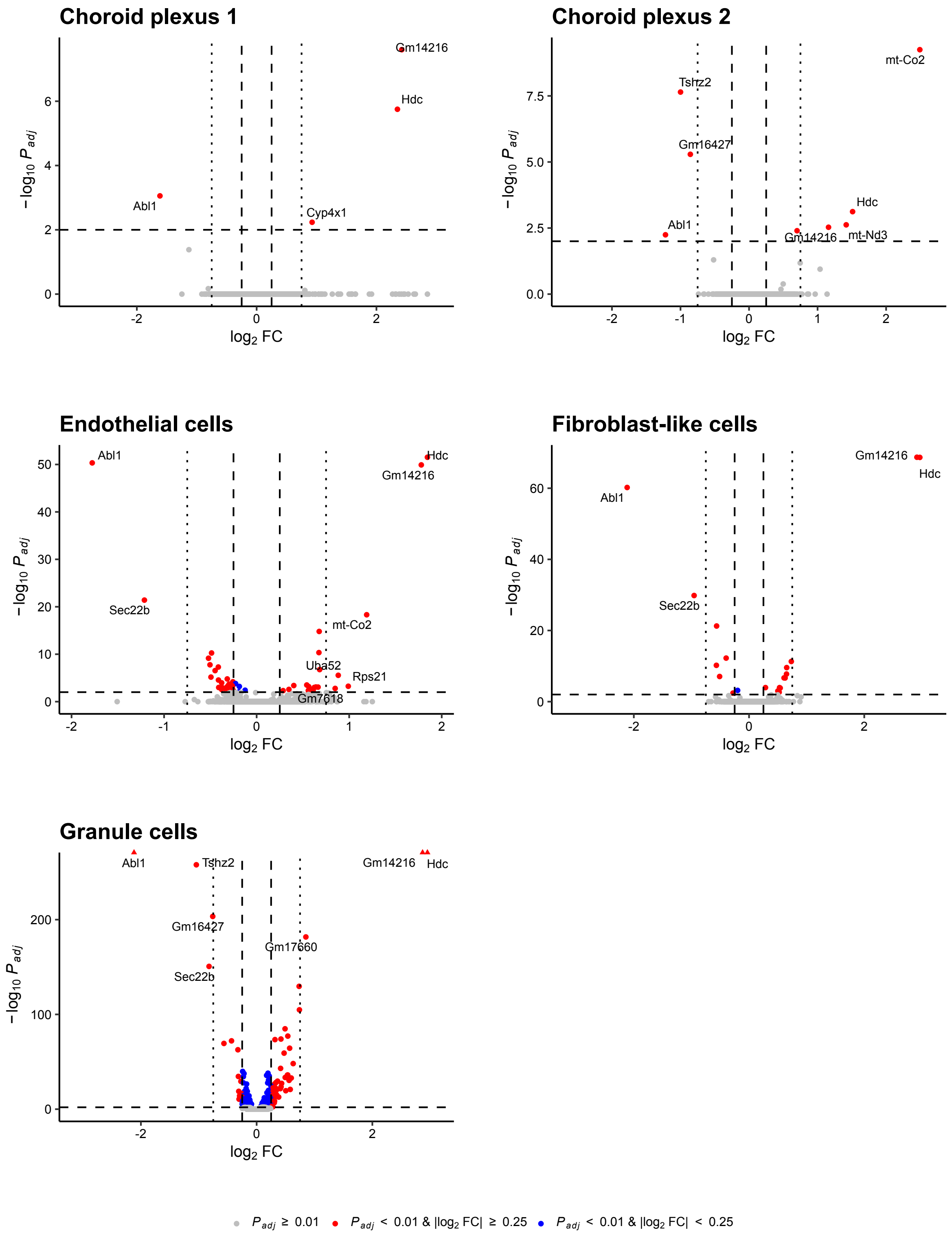


**Volcano plots resulting from comparison of *Pigv*$^{341E}$ cells and wild-type cells within other cellular subgroups.** Dashed horizontal lines are located at an adjusted p-value of 0.01, dashed vertical lines at an absolute $\log_2$ fold change of 0.25 and dotted vertical lines at an absolute $\log_2$ fold change of 0.75. For genes with an adjusted p-value below 0.01 and a minimum absolute $\log_2$ fold change of 0.75, gene symbols are depicted.

### Supplementary Material and Methods

#### Flow cytometry of mouse embryonic fibroblasts and mouse embryonic stem cells

Isolation of mouse embryonic fibroblasts (MEFs) from E13.5 embryos [Wt (n=4), het-*Pigv*$^{341E}$ (n=4), hom-*Pigv* (n=5)] is described elsewhere (McKean et al., 2012). MEFs were maintained in Dulbecco's modified eagle's medium (DMEM) supplemented with 10% fetal calf serum (FCS), 1% ultra-glutamine and 1% penicillin/streptomycin. mES cells were maintained as previously described (Franke et al., 2016) . For flow cytometry, cells were detached with cold ethylenediaminetetraacetic acid (EDTA) diluted in 1x phosphate buffered saline (PBS) and a cell scraper. Afterwards, cell suspension was pipetted several times to get a single cell suspension. Cells were stained with fluor-proaerolysin (FLAER) [Af-488] and Anti-mCD90/Thy1-PE (only used in mES cells) (R&D Systems, FAB7335P) in 1x PBS supplemented with 2% FCS for 30 min. Cells were then centrifuged at 400g for 5 min and washed in 1x PBS supplemented with 2% FCS two times. The stained cells were subsequently analyzed using the MACSQuant® VYB flow cytometer and the FlowJo™ v. 9.8.2 software. For mouse embryonic stem (mES) cells the average and standard deviation was calculated including two technical replicates.

References:

McKean DM, Niswander L. Defects in GPI biosynthesis perturb Cripto signaling during forebrain development in two new mouse models of holoprosencephaly. Biol Open. 2012;1(9):874–83.

Franke M, Ibrahim DM, Andrey G, Schwarzer W, Heinrich V, Schöpflin R, et al. Formation of new chromatin domains determines pathogenicity of genomic duplications. Nature [Internet]. 2016;538(7624):265–9. Available at: http://dx.doi.org/10.1038/nature19800

#### CRISPR-Cas9 gene-editing in mouse embryonic stem cells

mES cells were maintained as previously described (Franke et al. 2016). For knock-in of pathogenic mutations (e.g. Pigv c.1022C>A) in mES cells, single-stranded oligo-deoxynucleotides (ssODN) (60 pMol) were co-transfected with pX459 vector from Addgene containing the single-guided RNA (sgRNA) for e.g Pigv#1. Cloning of sgRNA in pX459 vector was performed according to Ran et al. (Ran et al., 2013). Transfection of mES cells and further processing was performed as previously described (Kraft et al., 2015). Genotyping of the cells were conducted as already explained (see Material and Methods, section: animals). Positive knock-in clones were confirmed by Sanger sequencing and used for diploid or tetraploid aggregation.

References:

Franke M, Ibrahim DM, Andrey G, Schwarzer W, Heinrich V, Schöpflin R, et al. Formation of new chromatin domains determines pathogenicity of genomic duplications. Nature. 2016;538(7624):265–9.

Ran FA, Hsu PD, Wright J, Agarwala V, Scott DA, Zhang F. Genome engineering using the CRISPR-Cas9 system. Nat Protoc. November 2013;8(11):2281–308.

Kraft K, Geuer S, Will AJ, Chan WL, Paliou C, Borschiwer M, et al. Deletions, inversions, duplications: Engineering of structural variants using CRISPR/Cas in mice. Cell Rep. 2015;10(5):833–9.

**Behavior testing**

Mice were habituated to the room adjacent to the testing room 30 min before starting behavior experiments. The equipment was cleaned with 70% ethanol before the experiments. In between animals and trials, the equipment was cleaned with 5% ethanol. Behavior experiments were conducted during the light phase (approx. 4 hours after the dark cycle ended). For the motor tests (rope grip, grip strength, beam walking, rotarod, foot print) nest construction test, home cage scan (HCS, CleverSys) recording and the social activity monitor (SAM, PhenoSys) (first approach) one cohort, Wt (female n=8, male n=4) *Pigv*$^{341E}$ (female n= 4, male n=2), was phenotyped at two consecutive time points (motor and nest construction test: 7 and 15 weeks, HCS: 8 and 16 weeks, SAM: 9 and 17 weeks). For cognitive tests (Barnes maze, y-maze), affective tests (open field, dark-light-box, elevated plus maze), social tests (three-chamber, social proximity), the marble burying tests and the buried food test two cohorts of the same age were used [first cohort: Wt (female n=4, male n=5) *Pigv*$^{341E}$ (female n=2, male n=2), second cohort: Wt (female n=5 male n=6) *Pigv*$^{341E}$ (female n= 2, male n=5)]. For the SAM (second approach) only the second cohort was used. The behavioral experiments were conducted in the same order in both cohorts. For the hindlimb clasping test a separate cohort was used [age: 6 weeks, Wt (female n=3, male n=5), het-*Pigv*$^{341E}$ (female n=4, male n=4), hom-*Pigv*$^{341E}$ (female n=4, male= n=6)].

**Hindlimb clasping test**

For each trial, mice were gently griped by the tale close to the body and held for 10 sec (3 trials in total). After each trial there was an (inter-trial interval) ITI of 5 min. During tail suspension, hindlimb clasping behavior was manually evaluated with following criterias: 0 - hindlimb were spread widely. 1 – one hindlimb pointed inwards for more than 50% of the time. 2 - both hindlimbs pointed inwards for more than 50% of the time in a repetitive manner. 3 - both hindlimbs were permanently pointing inwards for more than 50% of the time

**Rope grip test**

A 40 cm rope was tautly attached to a blue crate (40 x 30 x 26 cm). The blue crate was supplemented with padding on the bottom. Mice were gently griped by the tail close to the body and were allowed to grab the rope in the middle with both forepaws. Mice were then released and the time to reach one of the ends of the rope as well as the time to fall off the rope was measured. Mice were allowed to hang a maximum of 60 sec. The motoric behaviors were scored as followed: Score 0-Animal felt off. Score 1-Animal hang onto the string by one or both forepaws. Score 2-Animal hang onto the string by one or both forepaws and attempted to climb onto the string. Score 3-Animal hang onto the string by one or both forepaws plus one or both hindpaws. Score 4-Hang onto the string by forepaws and hindpaws plus tail was wrapped around string. Score-5-Animal escaped to the supports. Raw data acquisition was done manually by the experimenter.

**Grip strength test**

Each mouse was weighed before the test. Mice were gently griped by the tail close to the body and were gently lowered over the top of the bar that was connected to the TSE Grip Strength Meter apparatus. As soon as the mice attached the bar with both fore paws they were gently pulled away until they released the bar. Grip strength of only the forelimbs were recorded automatically and transferred to the computer by a (TSE) Grip Strength Meter. For each animal and type of measure, three values [g] were taken and averaged. Each animal of a batch was tested consecutively in the same order the grip strength mean value in [g] that was then normalized to the weight of the mice.

**Beam walking test**

Wooden beams (100 cm) were placed horizontally, 50 cm above the bench surface, with one end mounted on a narrow support and the other end attached to an enclosed box into which the mouse could escape. The ground under the beam was supplemented with padding. Mice were put on the bright lit start position and trained for two days to traverse a 20-mm diameter beam and enter an escape box on the other side of the beam. For training, 4 trials per animal were performed. On the test day, mice had to traverse the beam in two consecutive trials on each beam (20 mm, 15 mm, 10 mm) progressing from the widest to the narrowest beam. Mice were allowed up to 60 sec to traverse each beam. The time that mice needed to traverse the beam and to enter the escape box was measured. In addition, the number of hind foot slips were count. Raw data acquisition was done manually by the experimenter.

**Rotarod test**

Mice were put in the TSE Systems Rotarod apparatus. This apparatus 3 cm diameter rod that had ~1 mm horizontal grooves on the surface, which are designed to provide the mouse with better grip whilst running the task. The rod length was 30 cm, sub-divided with four partitions to create separate compartments such that five mice could be run simultaneously. Each mouse completed 3 trials with an ITI of 15 minutes. After the start of the test the rotarod accelerated then from 4-40 rpm for 300 sec. The latency to fall of the rotarod was measured by the TSE Rotarod software.

**Foot print test**

For this test mice were held in a scruffed position in order to apply color paints to their feet (red-forelimbs, blue-hindlimbs) (Dr. Oetker, Food colors). After the color was applied, for both training and testing, mice traverse the horizontal lane (67 cm long, 5 cm wide) of a T-maze covered with a white sheet, for two consecutive trials each. After the test, different parameters on the sheet were measured: stride length-forelimbs/hindlimb (SL-FL, SL-HL) (distance for forward movement for hindlimb and forelimb), hind base/fore base width (HBF, FBW) (distance between left and right footprints for hindlimb and forelimb), S-(distance between hindlimb and forelimb or overlap between forepaw and hindpaw placement). Raw data acquisition was done manually by the experimenter.

**Nest Construction Test**

Each Mouse was singly housed in a new home cage with fresh bedding that included a nestlet (5x5 cm pressed white cotton square). After 16 hours the nest built out of the cotton was scored manually with following criterias: Score 1-The nestlet was largely untouched (>90% intact). Score 2-The nestlet was partially torn up (50-90% remaining intact). Score 3-The nestlet was mostly shredded but often there was no identifiable nest site: < 50% of the nestlet remained intact but < 90% was within a quarter of the cage floor area, i.e. the cotton was not gathered into a nest but spread around the cage. Score 4- > 90% of the nestlet was torn up, the material was gathered into a nest within a quarter of the cage floor area, but the nest was flat with walls higher than mouse body height on less than 50% of its circumference. Score 5- > 90% of the nestlet was torn up, the nest was a crater, with walls higher than mouse body height on more than 50% of its circumference. Furthermore, the nestlet was weighed before the test and afterwards. The difference between both time points was calculated in [%].

**Home cage scan and computational analysis**

Mice were put in a home cage separated from each other and monitored for 23h. For the quantification of natural animal behavior in the familiar environment of its home cage, we used the HCS video and analytic software (CleverSys) which detects automatically different behaviors of freely moving mouse in its home cage (35 × 20 × 15 cm). Nineteen different behavior patterns of the mice were detected during a 23 h period (11 h of light phase and 12 h of dark phase/mouse). Number of occurrences and duration of each behavior were analyzed over time and on average per hour using custom-made R scripts.

**Social activity monitoring and computational analysis**

Mice were monitored for 14 days (first approach) with mixed genotypes in their home cage. For the second approach we monitored mice for 4 days with mixed and separated (non-mixed) genotypes. During this time the animal cages were placed on the SAM system (Phenosys. Germany). Each cage is placed on an ID-Grid sensor plate that detects the individual animals and tracks them as they move throughout the cage. Each ID-Grid sensor plate contains eight RFID sensors that records data on the location of the animals at every time point. The data is collected using Phenosoft Control software, preprocessed by Phenosoft analytics software and analyzed in R (R Core Team, 2017).

References:

R Core Team (2017). R: A language and environment for statistical computing. R Foundation for Statistical Computing, Vienna, Austria. URL https://www.R-project.org/.

**Barnes maze test**

During the training phase animals were given 4 trials a day for 4 days with an ITI of 15 minutes. For each trial the animals were placed on a table with a 92 cm diameter, containing 20 holes along the periphery in which one of the holes contained a nest that the animal could go into. The animal was allowed 3 minutes to find the nest. To encourage the animal to search for the nest a light white noise was played. Once the animal found the nest the white noise was stopped and the animal was allowed to stay in the nest for 1 minute. If the animal did not find the nest in the allotted 3 minutes it was shown the nest and the white noise was stopped. The animal was then allowed to stay in the nest for 1 minute. During the probe trials on day 5 and day 12 animals were returned to the table containing the 20 holes with the nest removed. During the probe trials the animal's behavior was observed for 90 s. For the evaluation of the search strategies 33.3% of the trials were analyzed for each animal. All parameters were recorded by the video-tracking system Viewer (Biobserve). Two animals had to be excluded from the analysis, since they were not able to learn the location of the exit hole during day 1-4.

**Buried food test**

For 3 consecutive days before testing, the mice were given sweet pellets (purified rodent tablets, 45 mg, TestDiet®, at least 3 times). All mice had to consume the pellets before the test or had to be excluded from the test. Four hours before testing, mice were deprived of food with water ad libitum. Before testing, each mouse was given 5 min of habituation in the testing cage (regular home cage; 35 × 20 × 15 cm) that contained 4 cm of fresh bedding. After this habituation, the mouse was temporarily removed from the testing cage, while a sweet pellet was hidden 1 cm deep and 5 cm away from the cage rear. For testing, the mouse was positioned in the center of the opposite end of the cage and the latency to find the food, i.e. the time from the moment the mouse was placed into the cage to the time it located the sweet pellet and initiated digging was recorded. Whether the pellet was eaten or not and other spontaneous behavior (rearing, climbing on the lid, grooming and extensive digging) were also scored. The mouse was then removed from the cage. A new cage and fresh bedding were used for each mouse. Raw data acquisition was done manually by the experimenter.

**Three-chamber Test**

The social testing arena was a rectangular, three-chamber box (60 x 40 x 22 cm, Stoelting apparatus) with white non transparent external walls. Each chamber was (20 x 40 x 22 cm) in size. Dividing walls were made from clear plexiglas, with rectangular openings (50 cm x 80 mm) allowing access into each chamber. The light condition was kept dim and uniform over the 3 chambers. In the experimental room, the test mouse was first placed in the middle chamber of the arena and allowed to explore for 10 min. To test sociability, after this habituation the mouse was trapped in the center chamber and an stranger mouse, that had no prior contact with the subject mouse, was placed in one of the side chambers. The location of the stranger mouse in the left vs. right side chamber was systematically alternated between testing animals. The stranger mouse was enclosed in a round wire cage with grey plexiglas covers (7 x 7 x 15 cm), which allowed nose contact through the bars but prevented fighting. The animals serving as stranger mice had previously been habituated to the small cage (at least 3 times before test, 5 min each time). An identical empty wire cage was placed in the opposite chamber. Both openings to the side chambers were then unblocked and the test mouse was allowed to explore the entire social arena for a 10 min session. The amount of time spent in each chamber and in close proximity to the grid were recorded by the video-tracking system Viewer (Biobserve). Number of sniffs at the wire cage was recorded manually as an index of close investigation. At the end of the first 10 min, each mouse was tested in a second 10 min session to quantify social recognition for a new stranger mouse after an ITI of 5 min (similar condition as the habituation phase). A second stranger mouse was placed into the previously empty wire cage. The testing mouse had a choice between the first, already-investigated mouse (familiar mouse=stranger 1), and the novel stranger mouse (stranger 2). As described above, measures were taken of the amount of time spent in each chamber and in close proximity with the grid during the second 10 min session. The position of stranger 1 and 2 was for each mouse changed in order to avoid side bias. Stranger mice had the same age and sex as the testing animal. One animal was excluded from the analysis since during habituation it only explored one side (chamber) of the maze.

**Y-maze**

Mice were placed in a y-maze that had 3 identical arms (34 x 6 x 17 cm), placed at 120° from each. The animals were placed facing the end of one arm of the maze. The starting arm was alternated between mice. Mice were allowed to explore the y-maze for 5 minutes. The percentage of spontaneous alternation (SAP), same arm returns (SAR), and alternate arm returns (AAR) as well as the distance traveled and time spent in each arm were recorded and measured using the video-tracking system Viewer (Biobserve).

**Marble burying test**

In a mouse cage (35 × 20 × 15 cm) twenty clean and identical glass marbles (green) were placed equally spaced in five rows of four marbles on a 4 cm layer bedding. For each animal a different cage with fresh bedding was used but illumination was kept homogeneous. The animal was placed in the testing cage close to a wall and allowed to explore the cage for 30 min. The latency for the first marble to be buried was manually scored. At the end of the test the mouse was rapidly removed from the cage and the total number of marbles buried was counted manually. A marble was considered "buried" when two thirds of its volume was covered by bedding. The marbles were cleaned with 5 % alcohol and dried for approximately 10 min before being used again.

**Social proximity test**

Mice were put in a transparent testing cage (size: 4x17 cm, 98cm$^2$) together with a stranger mouse with same age and sex. The behavior of the mice were recorded for 20 minutes. The video was manually evaluated after all experiments were finished. Number of different social traits were count: nose tip-to-nose-tip contact: The scored animal nose-tip and/or vibrissae contact with the nose-tip and/or vibrissae of the other animal. Nose-to-head contact: The scored animal nose-tip and/or vibrissae contact the dorsal, lateral, or ventral surface of the other animals head. Nose-to-anogenital contact: The scored animal nose-tip or vibrissae contact with the base of the tail and/or anus of the other animal. Crawl over: The scored animal forelimbs cross the midline of the dorsal surface of the other animal. Crawl under: The scored animal head goes under the ventral surface of the other animal. Rear up: Mice stretch their body vertically towards the cage. jump escape: the scored animal makes a vertical jump with all four feet leaving the ground.

**Dark/light Box**

Within the TSE multi-conditioning system mice were placed in the light chamber of a two compartment box that consisted of a light chamber and a dark chamber (30 x 22 cm for each chamber). Mice were then allowed to explore both chambers for a total of 10 minutes. The time that mice explored each the light and dark chambers  was measured and recorded by the TSE software.

**Elevated Plus Maze**

The mice were placed into a white PVC (50x50x53) elevated plus maze with two open arms and two closed arms and allowed to explore for 5 minutes.). The total distance, average velocity and the number of visits to each arm were recorded using the Viewer software (Biobserve).

**Open field test**

Mice were allowed to explore to the open field (50 x 50 cm) for 10 min. The time spent in the center and the periphery was recorded and measured by the Viewer software (Biobserve).

**Preparation of paraffin-embedded sections**

Animals were deeply anesthetized with Ketamin/Rompun and subsequently transcardially perfused with 20 ml 1x DPBS and with 20 ml paraformaldehyde (PFA) (4%) in 1x PBS with a 23G needle. The brain was removed and fixated in 4% PFA at room temperature over night. Thereafter, the brain was washed in 1x PBS twice for 30 min and finally in 50% and 70% Ethanol at room temperature for 30 min in each solution. The brain was stored in 70% ethanol at 4°C or further processed. Paraffin infiltration was performed with the tissue processor (Leica). Afterwards, paraffin-infiltrated brains were embedded in paraffin and sectioned (coronal) with a thickness of 5 microns using the paraffin microtome (Leica). Paraffin-embedded sections were deparaffinized in Xylene twice for 5 min. Afterwards, sections were rehydrated by washing them in 100% (twice), 90%, 70%, and 50% Ethanol for 5 min in each solution. Sections were then immediately processed for Nissl staining or synaptophysin-staining.

**Synaptophysin-staining**

Paraffin-embedded sections were rehydrated as described and then washed twice in 1× PBS for 3 min. Afterwards, antigen retrieval of sections was performed in 10 mM citrate buffer for 5 min at 60 °C, and the sections were washed twice in 1× PBS for 3 min. Sections were permeabilized and blocked for 30 min at room temperature in 0.2% Triton X-100 and 3% bovine serum albumin (BSA). Sections were washed twice for 3 min in 1× PBS and stained overnight at 4°C with anti-synaptophysin antibody (1:800, Abcam, ab32127), diluted in 1× PBS supplemented with 3% BSA. Afterwards, sections were washed for 1 h in 1× PBS, and secondary antibody (anti-rabbit-488Af, 1:1000) was applied in 1× PBS supplemented with 3% BSA. Sections were washed with 1× PBS for 3 min and stained with 4′,6-diamidino-2-phenylindole (DAPI) (1:1000) for 10 min. Finally, sections were washed three times for 3 min and embedded in Fluoromount.

For each biological replicate [Wt(female n=3, male n=2) Pigv341E(female n=2, male n=4), age: P23-25], three sections were analyzed. Three images were taken for each section per area [*Cornu Ammonis* 1 – *Stratum Radiatum* (CA1–SR), *Cornu Ammonis* 3 – *Stratum Radiatum* CA3–SR, *Cornu Ammonis* 1 – molecular layer of dentate gyrus (CA1–ML)] at 63× magnification on a ZEISS LSM 700 confocal microscope. 6 Z-stacks were recorded and transformed to one image with the ZEISS Zen Blue Software. Images were analyzed with FIJI ImageJ for quantification of synaptophysin immunoreactivity/µm2.

**Nissl staining**

Paraffin-embedded sections were deparaffinized and rehydrated as described above. Sections were then washed twice in dd$H_2O$ for 5 min. Afterwards, sections were maintained in Cresyl Violet buffered in Acetate Buffer (Nissl-staining solution) (pH 3,8-4) for 10 min. After Nissl-staining sections were washed twice for 3 sec in 100% Ethanol. Finally, the sections were embedded in Entellan. Images of sections were taken at 4x magnification.

**Acute slice preparation**

Acute hippocampal brain slices were prepared as described by Stempel et al. 2016. Adult mice were sacrificed by cervical dislocation. The brain was quickly removed and chilled in ice-cold sucrose - based artificial cerebrospinal fluid (sACSF) containing (in mM): NaCl 87,

NaHCO3 26, sucrose 50, glucose 10, KCl 2.5, NaH2PO4 1.25, MgCl2 3, CaCl2 0.5, continuously oxygenated. Horizontal slices (300 µm) were cut and stored submerged in sACSF for 30 min at 35 °C and subsequently stored in ACSF containing (in mM): NaCl 119, NaHCO3 26, glucose 10, KCl 2.5, NaH2PO4 1, CaCl2 2.5 and MgCl2 1.3 saturated with 95% (vol/vol) O2/5% (vol/vol) CO2, pH 7.4, at room temperature. Experiments were started 1 to 6 h after the preparation.

References:


Stempel AV, Stumpf A, Zhang HY, Özdoğan T, Pannasch U, Theis AK, et al. Cannabinoid Type 2 Receptors Mediate a Cell Type-Specific Plasticity in the Hippocampus. Neuron. 2016;90(4):795–809


**Schaffer collateral recordings**

Acute hippocampal brain slices were prepared as described above. Recordings were performed at room temperature (22–24°C) in a submerged recording chamber (Warner Instruments, RC-27L) ACSF, with solution exchange speed set to 2.5 ml/min. Low-resistance stimulation electrodes were placed in CA3–SR to stimulate Schaffer collaterals, and the recording electrode was placed in the CA1–SR field. Basal stimulation was applied every 10 sec. To analyze the input–output relationship, stimulation intensities were adjusted to different fiber volley (FV) amplitudes (0.05 mV increments) and correlated with the corresponding field excitatory post-synaptic potential (fEPSP) amplitudes. Paired pulse ratios (PPRs) were determined by dividing the amplitude of the second fEPSP (50 ms inter-stimulus interval) by the amplitude of the first (average of ten repetitions). Four high-frequency trains (100 pulses, 100 Hz) were applied every 10 sec, and the amplitude of the following fEPSP was normalized to the baseline [post-tetanic potentiation (PTP)]. Several measurements were conducted for each animal. The mean was calculated from all measurements from all animals [Wt(female n=2, male n=2) Pigv341E(female n=2, male n=2)].

**Pentylenetetrazol (PTZ) kindling model.**

Convulsive seizures were induced by repetitive intraperitoneal injections of PTZ every 10 min as described before (Van Loo et al., 2019) in Wt (female n=3, male n=9) and *Pigv*$^{341E}$ mice (female n=5, male n=6). Each injection consisted of 10 mg/kg PTZ (Sigma-Aldrich) and was given until a convulsive seizure occurred. The total dose of PTZ per animal did not exceed 100 mg/kg.

References:

Van Loo KMJ, Rummel CK, Pitsch J, Müller JA, Bikbaev AF, Martinez-Chavez E, et al. Calcium channel subunit α2σ4 is regulated by early growth response 1 and facilitates epileptogenesis. J Neurosci. 2019;39(17):3175–87.

**Isolation of hippocampal cells**

The animals were deeply anesthetized with Ketamin/Rompun and transcardially perfused with 20 ml 1x DPBS (pH 7,4) via a 20 ml syringe and a 23 G needle. The brains were removed and the hippocampi (2 hippocampi, each animal) were surgically taken out.

All procedures conducted for hippocampal cell isolation were performed on ice or at 4°C and all solutions were pre-cooled to 4°C. We pooled in total 8 hippocampi of 4 male animals per genotype (Wt and *Pigv*$^{341E}$). Hippocampi were dissociated in 1,5 ml Hibernate A (BrainBits) with a Dounce homogenizer (Active Motif) using the loose mortal. Single-cell suspension was transferred through a 70 µm strainer into a falcon. Afterwards, the Dounce homogenizer was washed twice with 1 ml Hibernate A. Remaining cells were collected from the washing step, passed through the 70 µm strainer and transferred into the falcon. Single-cell suspension was centrifuged at 500g for 10 min. Thereafter, pellet was resuspended in 1,55 ml DPBS (pH 7,4). Afterwards, 450 µl isotonic Percoll (10x DPBS diluted 1:10 in Percoll, pH 7,4) was added and single-cell suspension was pipetted up and down. 2 ml of 1x DPBS was very gently transferred on the single-cell suspension. The layered solution was then centrifuged for 10 min at 3250g. The supernatant with the myelin disk was removed immediately and the remaining pellet was washed in 4 ml 1x DPBS (pH 7,4) and centrifuged for 10 min at 400g. Finally, the pellet was resuspended in 1x DPBS (pH 7.4) and the volume was adjusted to get a final concentration of 10000 cells per 10 µl. Single-cell suspension was then further processed for single-cell RNA sequencing-library preparation.

**Single-cell RNA sequencing, preprocessing, and analysis**

Sequencing libraries were generated using the Chromium Single-cell Gene Expression Solution v3 (10X Genomics, Inc.) and sequenced on an Illumina NovaSeq 6000. Both pooled *Pigv*$^{341E}$ and wild-type samples were sequenced on the same lane. Demultiplexed sequencing data were provided by the core facility. Quality assurance of raw sequencing reads was performed with FastQC. For alignment to reference genome GRCm38.96 and quantification of single-cell gene expression, the zUMI pipeline with internal usage of the STAR aligner was employed. Both exonic and intronic reads were considered for downstream analysis.

Further processing and analysis of the generated gene expression count table were conducted in R, mainly using the single-cell genomics toolkit Seurat. Based on the distribution of cells, 19 out of 15,949 cells with ≤ 100 genes, ≤ 100 unique molecular identifiers (UMIs), or ≥ 30% of UMIs mapping to mitochondrial genes were removed by filtering (SI *Appendix*, Fig. S12). Gene expression values of the remaining 8,836 Pigv341E mutant cells and 7,094 wild-type cells were normalized using the “sctransform” procedure developed by Hafemeister and Satija (2019), including the percentage of UMIs mapping to mitochondrial genes as a regression variable (Hafemeister and Satija, 2019). After principal component analysis (PCA), the first 11 principal components were selected for dimensionality reduction based on the amount of captured variance (SI *Appendix*, Fig. S13). These components were used as input for clustering of cells, as well as for two-dimensional visualization with UMAP. For cluster definitions, we employed the unsupervised graph-based approach implemented in Seurat with k = 10 nearest neighbors. The resolution parameter was set to 0.2 based on visual concordance with the 2D representation in UMAP space. We then assigned cellular identities by analyzing the genes differentially expressed between the clusters. Candidate genes were compared with the adult mouse brain atlas (http://dropviz.org/) and in situ hybridization data from the Allen brain atlas (https://mouse.brain-map.org/search/index) (Saunders et al., 2018) (SI *Appendix*, Fig. S15). Distributions of cells across defined cellular subgroups were compared between genotypes by Pearson’s Chi-squared test. For differential expression testing between groups of cells, the default settings in Seurat were used. Genes with an absolute average log2(fold change) ≥ 0.25 and Bonferroni-adjusted p value ≤ 0.01 were considered significantly differentially expressed. Genes with significant differential expression between groups of interest were examined for overrepresentation of gene ontology (GO) terms relative to the background of all genes expressed in our data set using the clusterProfiler package; terms with an FDR adjusted p-value < 0.05 were considered significantly enriched.

References:

Hafemeister C, Satija R. Normalization and variance stabilization of single-cell RNA-seq data using regularized negative binomial regression. Genome Biol. 2019;20(1):1–15.

Saunders A, Macosko EZ, Wysoker A, Goldman M, Krienen FM, de Rivera H, et al. Molecular Diversity and Specializations among the Cells of the Adult Mouse Brain. Cell [Internet].2018;174(4):1015-1030.e16.Available at: https://doi.org/10.1016/j.cell.2018.07.028
